# Interactions between non-prion and prion domains of Rnq1 direct formation of amyloid vs liquid-like aggregates and create transmission barriers

**DOI:** 10.1101/2025.01.14.633072

**Authors:** Sangeun Park, Dania M. Maldonado, Michele L. Kadnar, Monica Andrade, Anna P. Fomitchova, Susan W. Liebman, Irina L. Derkatch

**Author notes:** Department of Biological and Vision Sciences, College of Optometry, State University of New York, New York, New York, United States of America (Monica Andrade). New York Medical College / New York Metropolitan Hospital, New York, New York, United States of America (Anna P. Fomitchova).

## Abstract

Prions are self-propagating protein conformations usually existing as amyloid aggregates. [*PIN^+^*], a prion form of the Rnq1 protein occasionally found in wild and laboratory yeast strains, facilitates both the *de novo* formation and destabilization of other yeast prions, and affects aggregation and toxicity of human misfolding disease proteins expressed in yeast. Rnq1 contains a short N-terminus with no confirmed function (the non-prion domain, NPD) and a C-terminus that carries four QN-rich regions and is sufficient for [*PIN^+^*] formation and maintenance (prion domain, PD). In the current study, a genetic screen identified the NPD T27P mutation that blocks transmission of the [*PIN^+^*] prion state from wild type Rnq1 (Rnq1_WT_) to mutant Rnq1_T27P_. The mutation doesn’t prevent Rnq1_T27P_ from switching to a prion state when overexpressed *in vivo*, or from forming amyloid fibers *in vitro*. Furthermore, like [*PIN^+^_WT_*], the newly formed [*PIN^+^_T27P_*]s promote the *de novo* appearance of the Sup35-based prion [*PSI^+^*]. We conclude that the NPD mutation creates a barrier for prion transmission from [*PIN^+^_WT_*] to Rnq1_T27P_. Because fluorescence microscopy shows that Rnq1_T27P_ efficiently joins [*PIN^+^_WT_*] aggregates, the barrier is likely due to the inability of Rnq1_T27P_ to propagate the specific [*PIN^+^_WT_*] conformational variant. Indeed, the analysis of [*PIN^+^_T27P_*]s resulting from rare transmission events from [*PIN^+^_WT_*] indicates that these [*PIN^+^_T27P_*]s must undergo conformational adaptation to yield more stable prion variants. Deletion analysis revealed that T27P constrains prion conformations through the first two QN-rich regions within the PD. The finding that Rnq1_T27P_-YFP readily forms non-amyloid liquid-like droplets, which Rnq1_WT_-YFP does not form, supports the idea that the NPD affects aggregation properties of the PD. We propose that these aggregation properties are essential for Rnq1’s functions, such as controlling aggregation of other proteins. This provides new insight into the role of heterologous proteins and transmission barriers in the origins of protein misfolding diseases.

**Author Summary:** Proteins must fold into the right shapes to work properly. Sometimes they fold incorrectly and stick together, forming long fiber aggregates that damage cells. This kind of “protein misfolding” causes human diseases such as Alzheimer’s. Certain yeast proteins behave similarly, making them useful to study this process. We investigate a yeast protein called Rnq1, which has a region that helps it misfold into fibers. These fibers can also cause other, unrelated proteins to misfold. We found that a mutation in a different part of Rnq1— outside the aggregation region — reduces the ability of non-mutant Rnq1 fibers to convert mutant Rnq1 into growing fiber aggregates. We also identified which section of the aggregation region is affected by this mutation. Interestingly, although the rarely converted mutant aggregates grow poorly at first, they can eventually “adapt” into a shape that grows better. The same mutation also pushes Rnq1 to form liquid-like droplets instead of fibers. Our findings show that the non-aggregating part of Rnq1 controls how Rnq1 aggregates, and, consequently, the appearance and elimination of aggregates formed by other proteins. Our work also helps explain how barriers to misfolded protein growth can be overcome, which is relevant to understanding human protein misfolding diseases.

## Introduction

The prion hypothesis postulates that some proteins can acquire stable unconventional conformations and pass these conformations to newly synthesized molecules with the same primary sequence. Amyloid, the β-sheet-rich fibers with protein molecules stacked on top of one another, is the structural basis for most prions and prion-like self-propagating protein conformations (Chuang et al., 2018; Dobson, 2017; Ke et al., 2020). Prions and amyloids attracted attention due to their connection to the so-called protein misfolding diseases that are associated with the formation of amyloid aggregates. The list of such diseases is expanding quickly from *bona fide* prion diseases, such as scrapie in sheep, mad cow disease in cattle, and Creutzfeldt-Jakob disease (CJD) in humans, which prompted the development of the paradigm of protein-based inheritance, to much more common neurodegenerative diseases, such as Alzheimer’s and Parkinson’s, as well as type 2 diabetes and, possibly, cancer (Ayers et al., 2020; Joshi and Ahuja, 2023).

Many protein misfolding diseases can be both sporadic, usually with a later age of onset, and hereditary, e.g. due to mutations in amyloid-forming proteins. This underscores the importance of uncovering the cellular mechanisms that drive and inhibit protein misfolding, as well as the control of conformational changes within specific aggregation-prone proteins that lead to disease-related aggregation. Prion diseases can also be infectious. Luckily, protective measures were stepped up in recent decades to prevent iatrogenic human to human transmission. Animal to human transmission is usually prevented by transmission barriers when differences in primary sequences of protein homologs from different species make interspecies transmission of the prion state inefficient (Beringue et al., 2008). However, as we learned from the appearance of a novel human disease resulting from a cross of the transmission barrier between cattle and humans, variant Creutzfeldt-Jakob disease (vCJD), such barriers may not be strong enough (Houston and Andreoletti, 2019). An important aspect of the problem is possible rapid adaptation of prion conformations in a new host after rare across-the-barrier transmission events (Weissman, 2012). Also, across-the-barrier transmission leads to forms of disease with distinct clinical manifestations compared to sporadic and hereditary diseases, like in the case of sporadic CJD and vCJD transmitted from cattle (Igel-Egalon et al., 2018).

Prion research has been greatly facilitated by the discovery of naturally occurring prions in the yeast *Saccharomyces cerevisiae* (Wickner, 1994; Liebman and Chernoff, 2012; Zhouravleva et al., 2023). The same underlying amyloid structure of yeast prions and human disease-related aggregation-prone proteins, as well as evolutionary conservation of genes makes yeast an excellent model for studying protein aggregation and mechanisms that control it. Many yeast prions carry long terminally located QN-rich domains that are unfolded in soluble proteins but are required and sufficient for taking on prion conformations and maintaining them. Hence their name – prion domains (PDs). Similar QN-rich or expanded polyQ domains also drive aggregation of some human disease proteins (Fuenteabla et al., 2010; Lieberman et al., 2019). [*PSI^+^*], the prion form of a translation termination factor Sup35 (eRF3), is the most extensively studied yeast prion, and many important findings come from the [*PSI^+^*] prion research (Liebman and Chernoff, 2012). One such finding is prion-prion interactions (Derkatch and Liebman, 2007). Initially, it was postulated that overexpression of a prion-forming protein should induce the *de novo* appearance of its respective prion conformation because increased protein concentration makes switching to a prion state more likely (Wickner, 1994). It was further postulated that if a prion is lost or cured from the cell, it should re-appear in the cell’s progeny either rarely spontaneously or frequently upon overexpression of the prion-forming protein (*ibid.*). Initial data for [*PSI^+^*] appeared to support these postulates (Derkatch et al., 1996). However, once [*PSI^+^*] was eliminated by the addition to yeast culture media of low concentrations of guanidine hydrochloride (GuHCl), a generic yeast prion curing agent, efficient induction of [*PSI^+^*] by Sup35 overexpression was no longer possible. This was because the treatment also cured the previously unknown [*PIN^+^*] prion required for induction of [*PSI^+^*] (Derkatch et al., 1997).

[*PIN^+^*] is formed by the Rnq1 protein (Sondheimer and Lindquist, 2000; Derkatch et al., 2001). Rnq1 consists of a short N-terminal domain with no confirmed function (the non-prion domain, NPD, aa 1-132; Fig 1A) and a C-terminal QN-rich domain sufficient for [*PIN^+^*] formation and maintenance (prion domain, PD; aa 132-405). The prion domain of Rnq1 is complex. It carries four discrete QN-rich regions. None of these alone is essential for [*PIN^+^*] prion maintenance, and any two are sufficient (Kadnar et al., 2010). Also, bacterially expressed purified Rnq1 fragments carrying NPD with only QN2, QN3 or QN4 form amyloid fibers *in vitro* (*ibid.*). The QN1 region is very short and does not form amyloid when attached to the Rnq1 NPD, but synthetic QN1 peptide does form amyloid fibers (*ibid*.). The [*mini-PIN^+^*]s formed by Rnq1 fragments with different combinations of QN-rich domains are all able to promote the *de novo* formation of [*PSI^+^*] upon Sup35 overproduction, *i.e.* no specific QN-rich region of Rnq1’s PD is essential for this prion-prion interaction (*ibid.*).

**Fig 1.**
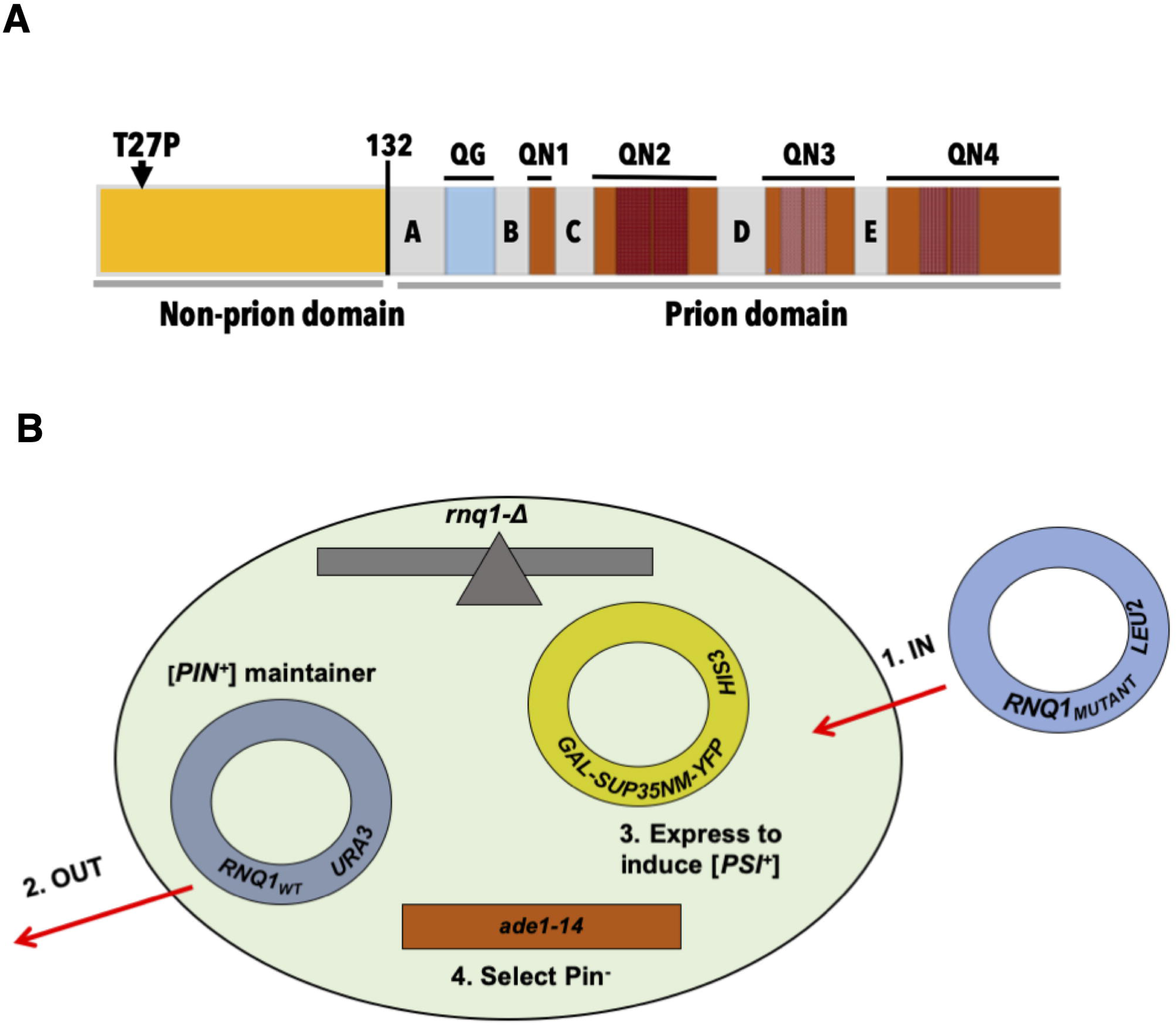
Genetic screen for mutations in Rnq1 that, when substituted for Rnq1_WT_ in a [*PIN^+^*] strain (Pin^+^ phenotype), lead to a Pin^-^ phenotype. (**A**) Structural organization of the Rnq1 protein. Shown are NPD (aa 1-132, yellow) with the position of the T27P mutation, and PD (aa 133-405) with a stretch of ten QG repeats (aa 153-172, blue), four distinct QN-regions (brown; QN1 through QN4), and five hydrophobic repeats with the LA^S^/_A_ ^L^/_M_ A core (light grey; A through E). The QN1 region is aa 185-198, QN2 - aa 218-263, QN3 - aa 279-319, QN4 - aa 337-405; patterned blocs indicate QN-rich repeats within QN2, QN3 and QN4). (**B)** Scheme of the genetic screen (see text).

Beyond promoting the *de novo* formation of [*PSI^+^*], [*PIN^+^*] is involved in other prion-prion interactions, both positive and negative. For example, [*PIN^+^*] also facilitates the *de novo* appearance of the [*URE3*] and [*SWI^+^*] prions with QN-rich PDs, even though the presence of [*PIN^+^*] is not a requirement (Bradley et al., 2002; Du et al., 2017), as well as some artificial prions (Borkhsenius et al., 2002) and the [HET-s]_Y_ prion formed in yeast by the *Podospora anserine* prion-forming protein Het-s with a non-QN-rich PD (Taneija et al., 2007). [*PIN^+^*] also interacts with non-prion aggregation-prone proteins, such as human Huntington’s and Machado-Joseph disease-associated proteins with expanded polyQ stretches (Osherovich and Weissman, 2001; Meriin et al., 2002) and TDP-43 associated with amyotrophic lateral sclerosis, frontotemporal dementia and limbic-predominant age-related encephalopathy (Park et al., 2017). An example of negative interactions is the mutual inhibition of stable inheritance by certain [*PSI^+^*] and [*PIN^+^*] prion variants: some weak [*PSI^+^*] variants are very unstable in the presence of some “single dot” [*PIN^+^*] variants, while some strong [*PSI^+^*] variants destabilize some “single dot” [*PIN^+^*]s (Bradley and Liebman, 2003; Mathur et al., 2009; Villali et al., 2020). Yet another type of interaction is a dramatic enhancement by [*PIN^+^*] of the [*SWI^+^*]-associated nonsense suppression in the strains lacking Sup35 PD (aka [*NSI^+^*], Nizhnikov et al., 2016).

Like other prions, [*PIN^+^*] depends on chaperone proteins for its maintenance. Some of these interactions are common with most other prions. For example, lack of the Hsp104 disaggregase cures [*PIN^+^*], as well as [*URE3*], [*PSI^+^*], and [*SWI^+^*] (Chernoff et al., 1995; Derkatch et al., 1997; Moriama et al., 2000; Du et al., 2008), likely because Hsp104 promotes conformational changes in prion aggregates and is needed to break growing prion aggregates to create smaller seeds that can be transmitted to daughter cells (Chernoff et al., 1995; Wegrzyn et al., 2001; Sapute-Krishnan et al., 2007). However, some prion-chaperone interactions are specific for a particular prion: the same chaperone can affect the formation of one prion but not others, and some chaperones have opposite effects on different prions (Barbitoff et al., 2022). [*PIN^+^*] has a highly specific interaction with Sis1, an Hsp40 co-chaperone of Hsp70 chaperones: not only does [*PIN^+^*] critically depend on Sis1 for maintenance (Sondheimer et al., 2001), but Sis1 is present in [*PIN^+^*] aggregates in equimolar amounts with Rnq1 (Lopez et al., 2003). In Sis1, the region critical for [*PIN^+^*] maintenance is within the G/F region that is not needed for the maintenance of other prions (Lopez et al., 2003). In Rnq1, the major site for interaction with Sis1 is located not in the PD, but in the NPD, in the LGKLALl sequence (Douglas et al., 2008).

Strikingly, most prion-forming proteins, including PrP in mammals and Rnq1, Sup35, and Ure2 in yeast, can form not just one but multiple prion variants (or prion strains) that differ in numerous prion-associated phenotypes (Bessen and Marsh, 1992; Bessen et al., 1995; Derkatch et al., 1996; Bradley et al., 2002; Schlumpberger et al., 2001). These prion variants manifest different conformations of prion-forming proteins in prion aggregates thus leading to differences in aggregate size, sensitivity to cellular chaperones and interactions with other prions *in vivo*, as well as in stability of corresponding amyloids *in vitro* (see above). Extensive studies of mutations within the Sup35 PD demonstrate that its primary sequence affects its conformational flexibility in terms of what [*PSI^+^*] strains the mutant protein can maintain (DePace, 1998; Chang et al., 2008; Bateman and Wickner, 2013).

Here we show that in Rnq1 the NPD is involved in determining the conformational flexibility of its PD and its ability to form particular amyloid variants and non-amyloid aggregates. Our findings are consistent with the hypothesis that the cellular function of the Rnq1 NPD is to regulate the aggregation of its PD and, through this aggregation, to control aggregation and disaggregation of other prions and prion-like aggregates.

## Results

### Genetic screen for *RNQ1* mutations leading to a Pin^-^ phenotype

Our goal was to investigate the structure-function organization of the Rnq1 protein in terms of its ability to maintain the [*PIN^+^*] prion and to facilitate the induction of the heterologous [*PSI^+^*] prion. Our approach was to screen for *RNQ1* mutant alleles (*RNQ1_MUTANT_*) that, when substituted for wild type *RNQ1 (RNQ1_WT_*) in a [*PIN^+^*] strain (Pin^+^ phenotype), lead to a Pin^-^ phenotype. The Pin phenotype was scored by examining the *de novo* appearance of the [*PSI^+^*] prion upon transient high-level expression of the PD of the [*PSI^+^*] prion forming protein Sup35 fused to YFP, Sup35NM-YFP. The Pin^+^ *vs* Pin^-^ phenotype was assigned, respectively, when [*PSI^+^*] appearance was detected *vs* not detected.

The screen is schematically shown in Fig 1B. **First**, *CEN LEU2* plasmids carrying PCR-mutagenized *RNQ1* driven by its native non-mutagenized promoter and followed by its native non-mutagenized terminator sequence were transformed into a [*PIN^+^*][*psi^-^*] strain where the chromosomal copy of *RNQ1* was deleted and [*PIN^+^*] was maintained by a similar *RNQ1_WT_* plasmid (*CEN URA3*). The strain also carried the *CEN HIS3* plasmid with the *SUP35NM-YFP* [*PSI^+^*] inducing construct controlled by the *GAL* promoter, and the *ade1-14* mutation allowing for [*PSI^+^*] detection. Transformants were selected on and then patched onto synthetic media selective for all plasmids, thus allowing for ∼20 generations of co-expression of Rnq1_WT_ and Rnq1_MUTANT_. **Second**, after two passages on media selective for only the mutagenized *RNQ1* and [*PSI^+^*] inducing plasmids, the *RNQ1_WT_* maintainer plasmid was shuffled out on media supplemented with 5-FOA and selective for the other two plasmids. **Third**, [*PSI^+^*] was induced by high-level expression of Sup35NM-YFP on SGal-LeuHis media. **Fourth**, cells carrying [*PSI^+^*] were selected on non-inducing media lacking adenine and not selective for plasmids. Selection of [*PSI^+^*] cells on adenineless media is based on nonsense suppression of the *ade1-*14 (UGA) mutation due to the recruitment of the Sup35 protein, a translation termination factor, into the [*PSI^+^*] prion aggregates. For Pin^+^ cultures, thick, even if heterogeneous, growth was observed over the entire patch. For the presumptive Pin^-^ cultures, either no growth or rare Ade^+^ colonies were observed on media lacking adenine. In such cultures, the Pin^-^ phenotype was confirmed by colony purifying yeast from the inducing SGal-LeuHis media onto YPD, where [*PSI^+^*] Ade^+^ cells form pink or white colonies that are clearly distinct from red Ade^-^ colonies formed by [*psi^-^*] cells. Finally, the Pin^-^ phenotype was confirmed by fluorescence microscopy of cells grown on the [*PSI^+^*] inducing galactose media using the YFP-tagged Sup35NM as a reporter for the lack of Sup35 aggregation. In Pin^+^ cultures the appearance of [*PSI^+^*] is visualized as ring- or dot-like foci, while in [*psi^-^*] cells Sup35NM fluorescence is evenly distributed (Patino and Lindquist, 1996; Zhou et al., 2001). To confirm that the Pin^-^ phenotype was due to a mutation in the shuffled-in *RNQ1* gene, *RNQ1_MUTANT_* plasmids were isolated from each candidate mutant, individually re-transformed back into the yeast strain used for the original screen, and tested according to the same scheme (see Materials and Methods for additional approaches used to improve the candidate pool).

Approximately 12,000 yeast transformants were screened and 54 were either firmly Pin^-^, or their Pin phenotype was dramatically reduced. 53 of these carried plasmids with either nonsense or frameshift *RNQ1* mutations. All nonsense mutations and most frameshifts were located in the N-terminal part of *RNQ1*, either within its NPD, or before the end of the second QN-rich domain within the PD (QN2, see Fig 1A). All five unique frameshift mutations that were located downstream of QN2 led to long C-terminal extensions due to out-of-frame translation. The only missense mutant that passed the screen carried the T27P (ACA ⇨ CCA) mutation in the beginning of the Rnq1 NPD (Fig 1A; also see Fig 2B).

**Fig 2.**
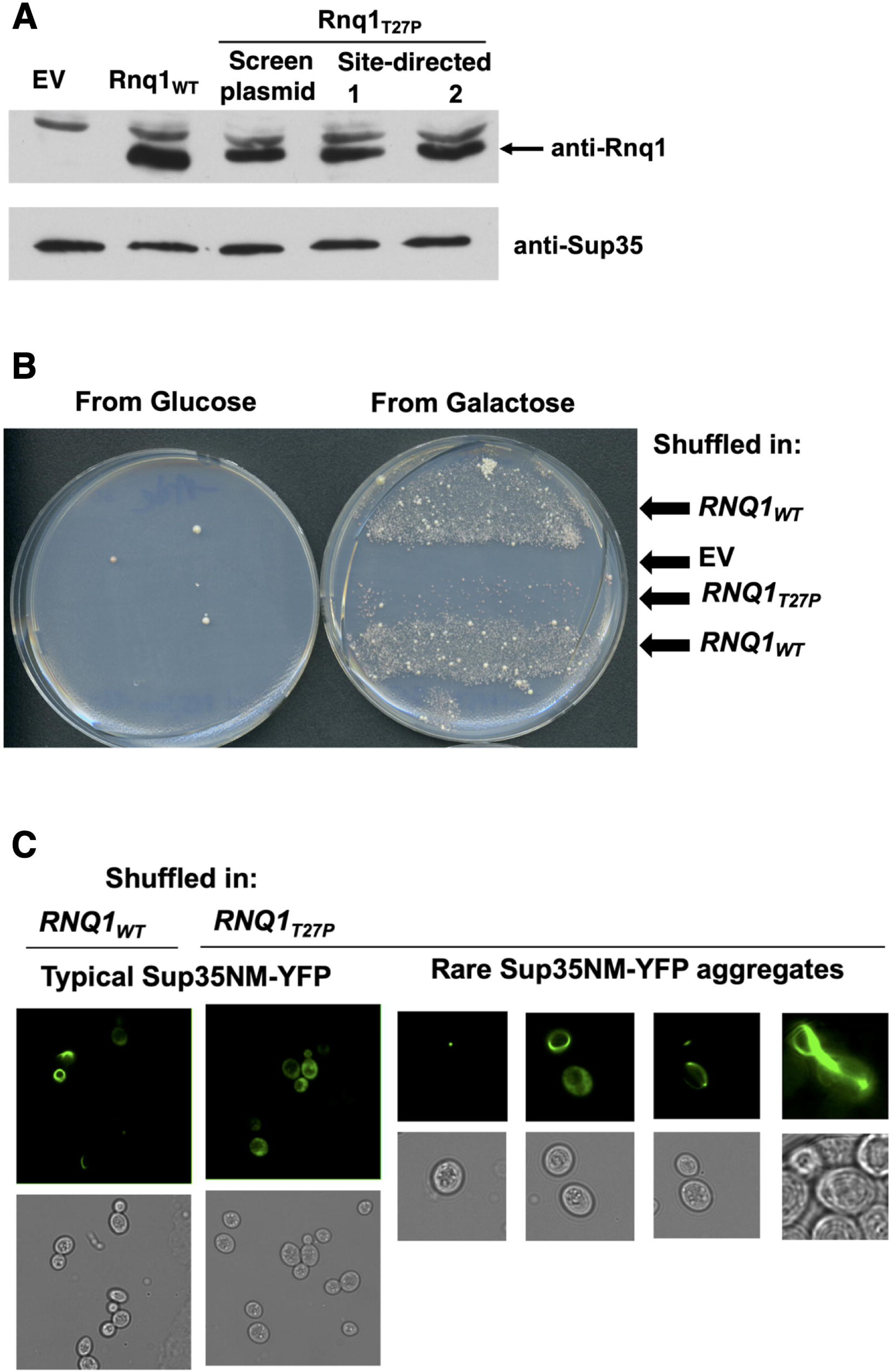
The T27P mutation in Rnq1 NPD leads to a dramatic reduction of the Pin^+^ phenotype when Rnq1_T27P_ is substituted for Rnq1_WT_ in a [*PIN^+^*] strain. (**A**) Western blot analysis of the expression of Rnq1_T27P_ and Rnq1_T27P_ in *the rnq1-Δ* strain. The *rnq1-Δ* YID193 strain was transformed with the indicated *CEN LEU2* plasmids. For *RNQ1_WT_* the pID129 plasmid was used. For*RNQ1_T27P_*, shown is expression from the original plasmid from the mutagenized library and from two independent isolates of the pID335 plasmid where the T27P mutation was introduced by site-directed mutagenesis. Expression of Sup35 is used as a loading control. Both panels are from the same Western blot: anti-Rnq1 and anti-Sup35 antibodies were added simultaneously. Mobilities of Rnq1 and Sup35 are as expected based on protein molecular weights. Expression of Rnq1_T27P_ is ∼2-fold lower compared to Rnq1_WT_. (**B)** Rnq1_T27P_ leads to a dramatic reduction of the Pin^+^ phenotype when substituted for Rnq1_WT_ in a [*PIN^+^*] strain. Shown is the image of the YID146.3 cells with *RNQ1_T27P_* (or indicated control plasmids) substituted for *RNQ1_WT_* incubated on SD-Ade media for 14 days at 30^0^C after previously being grown either on the [*PSI^+^*] inducing SGal-LeuHis media, or on the control SD-LeuHis (two passages). Each row includes patches of 5 independent transformants. See Fig S1 for images of the same cultures on SD-LeuHis and and SGal-LeuHis from which cells were transferred onto SD-Ade. (**C**) Upon high-level expression of Sup35NM-YFP in the cultures where *RNQ1_T27P_* is substituted for *RNQ1_WT_*, the *de novo* forming Sup35 aggregates are rare, but similar in appearance to aggregates forming in the control *RNQ1_WT_* cultures. Shown are fluorescent and bright-field images of cells carrying *GAL-SUP35NM-YFP* and the indicated *RNQ1* construct grown on SGal-LH for ∼48hrs at 30^0^C after the plasmid shuffle described in Fig 1B. See Table 1 for quantification. A typical group of cells is shown for the *RNQ1_WT_* culture. For *RNQ1_T27P_*, all cells have the evenly distributed fluorescence in a typical group of cells representative of the 97.5% of the culture. Additionally shown for *RNQ1_T27P_* are rare cells with dot-, line-, and ring- aggregates, as well as aggregate-containing mother-daughter pairs (a line and a dot, or a ring shared by a mother and a daughter).

**Table 1.** Analysis of Sup35NM-YFP aggregation in *RNQ1_T27P_* cells after shuffling out the *RNQ1_WT_* plasmid indicates low-level [*PSI^+^*] induction. Aggregation of Sup35NM-YFP was analyzed in YID146 cells after substituting the *RNQ1_T27P_ LEU CEN* or indicated control constructs for *URA3 CEN RNQ1_WT_* (see Fig 1B and text). To induce the expression of Sup35NM-YFP, yeast passaged on FOA media to eliminate the *URA3 CEN RNQ1_WT_* plasmid were replica plated onto the SD-LeuHis and then SGal-LeuHis media and incubated for ∼48 hours at 30^0^C. For *RNQ1_WT_* and *RNQ1_T27P_*, at least 11 vision fields were scored for each construct, ∼500 cells total per construct. For EV, no aggregates were detected in ∼1,000 cells scored. Based on T-test, differences are statistically significant between Rnq1_T27P_ *vs* Rnq1_WT_ (p<10^-13^), Rnq1_T27P_ *vs* EV (p<0.005), and Rnq1_WT_ *vs* EV (p<10^-14^). Qualitative assessment of aggregation in >10,000 cells in the cultures used for quantification further confirmed the difference between the *RNQ1_T27P_* and EV: bright ring- or dot-aggregates were readily detected in Rnq1_T27P_ expressing cells, but no ring aggregates were detected in EV cells, where there were only a few dot foci, mostly atypically large. Also, similar observations were made upon qualitative assessments after a shorter (overnight) incubation on SGal-LeuHis and after two passes on this medium for over 4 days.

| Construct after plasmid shuffle | % cells with Sup35NM-YFP aggregates |  |  |
| --- | --- | --- | --- |
|  | Average | SD | SEM |
| $RNQ1_{WT}$ | 56.8 | $\pm 14.5$ | $\pm 3.0$ |
| $RNQ1_{T27P}$ | 2.6 | $\pm 3.1$ | $\pm 0.8$ |
| EV | 0 | NA | NA |

Identification in the screen of constructs synthesizing only the NPD or part of it was expected because of the requirement of the PD of Rnq1 for the maintenance of the [*PIN^+^*] prion (Vitrenko et al., 2007a; Kadnar et al., 2010). The fact that no nonsense mutations were detected beyond the second QN-rich region can be explained based on our earlier deletion analysis of Rnq1 (Kadnar et al., 2010): the QN-rich regions within Rnq1 PD are redundant in their ability to acquire the [*PIN^+^*] prion state from Rnq1_WT_ and maintain the Pin^+^ phenotype (i.e. allow for [*PSI^+^*] induction). Specifically, the N-terminal Rnq1 fragments including at least QN1 and entire QN2 were sufficient to maintain Pin^+^, so mutants retaining these regions were not expected to pass the screen. However, deletion analysis also indicated that the C-terminal location of the QN2 region in the truncated construct carrying only QN1 and QN2 was important for the transfer of [*PIN^+^*] from Rnq1_WT_ (*ibid*), so the identification of frameshifts leading to long extensions downstream of QN2 is also not surprising. Finally, the fact that we have not identified PD-located missense mutations can also be explained by the structural redundancy of the Rnq1 PD: a single mutation that would just inactivate one of the four QN-rich regions is unlikely to block [*PIN^+^*] propagation or the ability of the mutant to promote [*PSI^+^*] induction.

### The inability to induce nonsense suppression in cultures where Rnq1T27P was substituted for Rnq1WT is due to the dramatic reduction in the ability to induce [*PSI^+^*]

For all experiments described here and below, we used plasmids with the T27P mutation introduced by site-directed mutagenesis, to exclude the possibility that additional mutations in the regions we did not sequence outside of the *RNQ1* gene may have contributed to the Pin^-^phenotype during the screen. We also reasonably excluded the possibility that the absence of [*PIN^+^_T27P_*] in most post-shuffle Rnq1_T27P_ cells was due to insufficient Rnq1_T27P_. Although there is an ∼2-fold difference in protein abundance between Rnq1_T27P_ and Rnq1_WT_ (Fig 2A) this is unlikely to be the main reason for the prion loss because (i) [*PIN^+^_WT_*] is 100% stable in diploids heterozygous for *RNQ1* deletions (Derkatch et al., 2001) and (ii) a similar reduction in abundance compared to full-length Rnq1_WT_ does not destabilize mini-[*PIN^+^*]s maintained by Rnq1 fragments (Kadnar et al., 2010).

The goal for further experiments was to understand what leads to the loss of the Pin^+^ phenotype of the *rnq1-Δ* yeast cultures expressing Rnq1_T27P_ after the elimination of Rnq1_WT_. As can be seen on Fig 2B, the cultures expressing Rnq1_T27P_ were almost Pin^-^ compared to the positive control expressing Rnq1_WT_. However, compared to the negative control where Rnq1_WT_ was substituted with the empty vector (EV), there were occasional Ade^+^ colonies on adenineless media on the Rnq1_T27P_ patches. Out of 24 such Ade^+^ microcolonies from six independent Rnq1_T27P_ transformants, 23 were [*PSI^+^*] according to the GuHCl test. The fluorescence analysis of Ade^+^ cultures that retained the *GAL-SUP35NM-YFP* plasmid revealed dot-like foci in most fluorescent cells in all eight cultures examined, and these foci were indistinguishable from [*PSI^+^*] foci in the control Rnq1_WT_ cultures. Thus, the Ade^+^ colonies in the cells expressing Rnq1_T27P_ after shuffle were due to [*PSI^+^*]s, so [*PSI^+^*] induction was not blocked completely due to the T27P mutation.

We then asked whether the [*PSI^+^*] induction was indeed ∼50-100-fold less frequent in the Rnq1_T27P_ *vs* the Rnq1_WT_ cultures, as it appeared, or alternatively it was close to normal levels but most newly forming [*PSI^+^*]s were not detected in the assay based on the ability of [*PSI^+^*] cells to grow on adenineless media. This could happen for five reasons. (1) Most [*PSI^+^*] variants forming in cells carrying [*PIN^+^_T27P_*] are “very weak” and unable to ensure robust growth on adenineless media. Indeed, some [*PIN^+^*] variants promote induction of predominantly weak [*PSI^+^*]s (Sharma and Liebman, 2013). (2) Most [*PSI^+^*]s promoted by [*PIN^+^_T27P_*] are highly unstable and are lost during the first few cell divisions. Indeed, some [*PSI^+^*] variants are permanently unstable (Sharma and Liebman, 2012), and some [*PIN^+^*] variants are known to destabilize weak [*PSI^+^*]s they induce (Bradley and Liebman, 2003). (3) In [*PIN^+^_T27P_*] cells, most newly forming Sup35 aggregates do not mature into [*PSI^+^*] or are lost in the absence of Sup35 overexpression. Such [*PSI^+^*] variants have been described by Salnikova et al. (2005). (4) Most [*PSI^+^*]s in [*PIN^+^_T27P_*] cultures are extremely toxic and lead to cell death on the Sup35NM-YFP inducing media, prior to transfer to adenineless media where Sup35NM-YFP is no longer expressed. Indeed, growth of [*PSI^+^*] cells is inhibited by Sup35 overexpression in a [*PSI^+^*] variant-specific manner (Derkatch et al., 1996). (5) Most [*PSI^+^*]s induced in [*PIN^+^_T27P_*] are toxic even in the absence of Sup35NM-YFP expression. Such “killer” [*PSI^+^*]s have also been described (McGlinchey et al., 2011). Experiments discussed below essentially exclude all these possibilities.

To assess the strength of [*PSI^+^*] variants induced in cells expressing Rnq1_WT_ and Rnq1_T27P_, yeast scraped up from patches incubated on SD-Ade media at 20^0^C for ∼25 days after [*PSI^+^*] induction were colony purified on YPD, and individual colonies were randomly picked for further analysis. Incubation at low temperature promotes nonsense suppression in weaker [*PSI^+^*] variants (Derkatch et al., 1996) and prolonged incubation, in our experience, allows even the weakest [*PSI^+^*]s to form microcolonies. We found that similar arrays of [*PSI^+^*] variants were induced in the presence of Rnq1_WT_ and Rnq1_T27P_, with no prevalence for weak [*PSI^+^*]s in the Rnq1_T27P_ cultures. Specifically, moderate [*PSI^+^*]s constituted 85% and 91%, weak [*PSI^+^*]s – 6% and 6%, and strong [*PSI^+^*]s – 10% and 3% for Rnq1_WT_ and Rnq1_T27P_, respectively (Table S1). The slightly smaller percentage of strong [*PSI^+^*] variants induced in Rnq1_T27P_ cells was not statistically significant (p>0.1) and could not explain the large difference in the Pin phenotype.

Furthermore, the dramatically reduced Pin^+^ phenotype in the Rnq1_T27P_ after-shuffle cultures was not due to extremely weak (essentially Ade^-^) or highly unstable [*PSI^+^*] variants. To test this, yeast were colony purified directly from the [*PSI^+^*] inducing galactose media onto YPD. Only ∼0.7% of >5,000 colonies originating from 12 independent *RNQ1_T27P_* transformants were white or pink, and most of these colonies were [*PSI^+^*] based on the GuHCl test and, also, the Sup35 aggregation test for those that retained the *SUP35NM-YFP* plasmid. The color of pink colonies and their ratio to white colonies was similar to that in the positive control cultures with *RNQ1_WT_* shuffled-in where white and pink (and [*PSI^+^*]) colonies constituted ∼20%. In the negative control cultures with EV shuffled-in, the rare (<0.1%) non-red colonies were usually white and not [*PSI^+^*]. This is consistent with results shown in Fig 2B. Importantly, no dark pink colonies indicative of very weak [*PSI^+^*] were detected in Rnq1_T27P_ cultures, and analysis of all slightly lighter shade of red colonies (∼30) showed that they were not [*PSI^+^*]. In addition, fluorescence microscopy of ∼40 typical red colonies retaining *GAL-SUP35NM-YFP* did not reveal any [*PSI^+^*]s. Finally, no increase of the proportion of sectored white/red or pink/red [*PSI^+^*] colonies was detected for Rnq1_T27P_ compared to Rnq1_WT_. The fact that newly forming [*PSI^+^*]s are frequently unstable is well-known, so the presence of some sectored colonies was expected (Sharma and Liebman, 2012). All potentially sectored colonies in Rnq1_T27P_ cultures yielded stable [*PSI^+^*]s upon further analysis. Altogether, these experiments exclude possibilities (1) and (2) from the list above.

Fluorescence microscopy of aggregate formation upon high expression of Sup35NM-YFP was used to record the initial steps of [*PSI^+^*] formation. In the after-shuffle Rnq1_T27P_ cultures only ∼2.5% cells had aggregates (Table 1; Fig 2C). This is significantly less compared to Rnq1_WT_ (∼57%) and significantly more compared to EV (<0.01%). Both ring-, line- and dot-shaped foci were detected in the *RNQ1_T27P_* carrying cells, and these foci were not distinguishable from those in cells with *RNQ1_WT_*. The presence of dot-like foci indicative of mature [*PSI*^+^] and occasional mother-daughter pairs with aggregates in both cells indicated that newly forming Sup35 aggregates were heritable in the Rnq1_T27P_ expressing cells. Also, there was no indication of increased cell toxicity in cells co-expressing Rnq1_T27P_ and Sup35NM-YFP: no abnormally shaped elongated cells, no increase in the proportion of non-fluorescent cells, and no growth inhibition compared to the Rnq1_WT_ control (Fig S1). Finally, in the [*PSI^+^*]s induced in the Rnq1_T27P_ expressing cells, maintenance of the *LEU2 RNQ1_T27P_* and *HIS3 GAL-SUP35NM-YFP* plasmids was not reduced compared to the maintenance of *LEU2 RNQ1_WT_* and *HIS3 GAL-SUP35NM-YFP* in the [*PSI^+^*]s induced in the Rnq1_WT_ cells (see description of [*PSI^+^*] isolation above and Table S1). Together, these experiments exclude possibilities (3), (4), and (5) above.

In summary, the cultures expressing Rnq1_T27P_ have a dramatically reduced but otherwise typical Pin^+^ phenotype.

### Overexpression of Rnq1T27P induces a heritable Pin^+^ phenotype indicative of the formation of [*PIN^+^T27P*]

In general, the *de novo* appearance of prions can be induced by their transient overexpression, and the process of prion aggregate formation can be monitored by microscopy by attaching a fluorescent tag to a prion forming protein (Zhou et al., 2001). However, Rnq1_WT_ does not form *de novo* [*PIN^+^*] aggregates upon overexpression when attached to a C-terminal GFP family tag (Derkatch et al., 2001), so a negative result for Rnq1_T27P_ would not be informative. Thus, the *de novo* induction of [*PIN^+^_T27P_*] and [*PIN^+^_WT_*] was attempted by overexpressing, respectively, the **untagged** Rnq1_T27P_ and Rnq1_WT_ driven by the inducible *CUP* promoter and scoring for the Pin^+^ phenotype by [*PSI^+^*] induction upon high-level expression of Sup35NM-YFP.

For the first experiment (Fig 3A), the *CEN URA3* plasmids carrying either the *CUP-RNQ1_T27P_* or *CUP-RNQ1_WT_* inserts, or just the empty *CEN URA3* vector, were introduced into a [*pin^-^*][*psi^-^*] *rnq1-Δ* strain already carrying the *HIS3 GAL-SUP3NM-YFP* [*PSI^+^*] inducing construct. Transformants were patched on SD-UraHis media where plasmid-borne constructs were not expressed. Then yeast cultures were replica plated onto the SGal-UraHis supplemented with 50μM CuSO_4_ where Rnq1_T27P_ or Rnq1_WT_ were overexpressed together with Sup35NM-YFP, or on control media where neither or only one of the two constructs was expressed (SD-UraHis, SD-UraHis+50μM CuSO_4_, and SGal-UraHis). Induction of [*PSI^+^*] was scored on adenineless media to which yeast were replica plated from either SGal-UraHis+50μM CuSO_4_ or control media. Indeed, heterogeneous growth on adenineless media indicative of [*PSI^+^*] induction was observed following co-overexpression of Sup35NM-YFP with either Rnq1_T27P_ or Rnq1_WT_, with no obvious effect of the T27P mutation on the efficiency of [*PSI^+^*] induction (Fig 3A). The *de novo* formation of [*PSI^+^*] was also confirmed by fluorescent microscopy: in cells from the SGal-UraHis+50μM CuSO_4_ media we observed ring-, line- and dot-shaped Sup35NM-YFP foci indistinguishable in appearance between the Rnq1_T27P_ and Rnq1_WT_ expressing cultures (data not shown). The limitation of this experiment was that it did not allow us to test if the Pin^+^ phenotypes induced by the overexpression of Rnq1_T27P_ and Rnq1_WT_ were heritable, indicative of the formation of, respectively, [*PIN_T27P_^+^*] and [*PIN^+^_WT_*], or only non-propagating aggregates were formed by one or both constructs.

**Fig 3.**
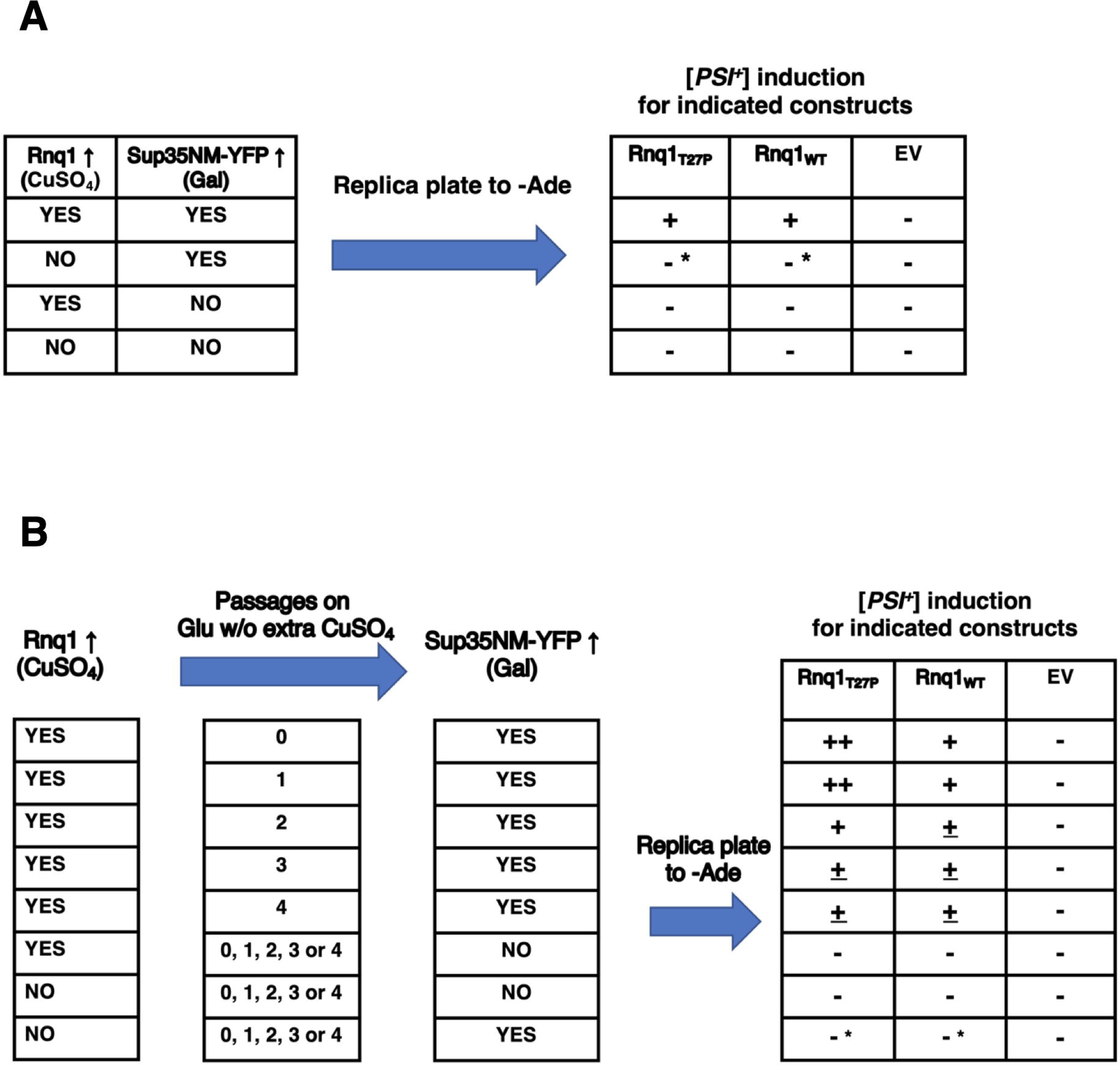
The T27P mutation does not interfere with the ability of the Rnq1 protein to acquire a heritable Pin^+^ state upon overexpression, indicative of the *de novo* formation of the [*PIN^+^_T27P_*] prion. For the description of Experiment 1 (**A**) and Experiment 2 (**B**) see Results and Materials and Methods. The (++), (+), and (±) indicate the relative level of [*PSI^+^*] induction based on growth on adenineless media compared to lack of [*PSI^+^*] induction (-), with (++) being the most robust. In (B) some negative control experiments are grouped together. * Occasional Ade^+^ colonies were detected after induction of the *GAL* promoter-driven Sup35NM-YFP in cultures where *CUP1*-driven *RNQ1_T27P (or WT)_* constructs were not overexpressed. This is indicative of a low-level of induction of Rnq1-based prions, probably due to residual expression of *CUP1-RNQ1* constructs in all or some cells on regular SD media which contains 0.16μM CuSO_4_. The number of such Ade^+^ colonies gradually increased after multiple passages prior to the induction of Sup35NM-YFP but never reached the level of Pin^+^ phenotypes observed in cultures that went through transient Rnq1_T27P (or WT)_ overexpression on 50μM CuSO_4_.

For the second experiment (Fig 3B), the [*pin^-^*][*psi^-^*] *rnq1-Δ* strain carrying the *HIS3 CEN GAL-SUP3NM-YFP* plasmid was co-transformed with *URA3 CEN CUP-RNQ1_T27P_* (to be the inducer of the [*PIN^+^_T27P_*] prion state) and the *LEU2 CEN* plasmid carrying *RNQ1_T27P_* under its original promoter (to be the maintainer of this prion state after induction). Control samples were co-transformed with corresponding *RNQ1_WT_* plasmids (*URA3 CEN CUP-RNQ1_WT_* and *LEU2 CEN RNQ1_WT_*), or with empty vectors (*URA3 CEN* and *LEU2 CEN*). The Pin phenotype was scored for by growth on adenineless media after overexpression of Rnq1_T27P (or WT)_ and *GAL-SUP3NM-YFP* induction but with 0 (co-overexpression), 1, 2, 3, or 4 passages on the non-inducing glucose no-copper plasmid-selective media between the Rnq1_T27P (or WT)_ overexpression and *GAL-SUP3NM-YFP* induction. The efficiency of [*PSI^+^*] induction was slightly but reproducibly higher in the Rnq1_T27P_ expressing cells compared to Rnq1_WT_ in the nonsense suppression test for most of ∼50 co-transformants tested for each set of plasmids, but no statistically significant difference was detected when aggregation was assessed by fluorescent microscopy (not shown). As can be seen in Fig 3B, the Pin^+^ phenotype of both Rnq1_T27P_ and Rnq1_WT_ expressing cultures became slightly weaker upon subsequent passages on media not supplemented with CuSO_4_. This is expected considering that newly forming prions are frequently unstable (shown before for spontaneous [*PIN^+^_WT_*]s by Derkatch et al. (2000) and newly induced [*PSI^+^*]s by Zhou et al. (2001)). Important is the retention of the robust Pin^+^ phenotypes after ∼30 generations (4 passages) on the no-copper media where Rnq1_T27P_ and Rnq1_WT_ were no longer overexpressed, which is indicative of the formation of heritable [*PIN_T27P_^+^*] and [*PIN^+^_WT_*] prions (for comparison, growth for a similar number of generations on 5mM GuHCl media that inhibits prion propagation is sufficient to completely cure [*PIN^+^*]). Also important is that Rnq1_T27P_ is not deficient in acquiring such a heritable [*PSI^+^*]-inducing state compared to Rnq1_WT_.

### Rnq1T27P forms amyloid fibers *in vitro*

Another way to assess the aggregation properties of Rnq1_T27P_ was to test the ability of the bacterially expressed and purified protein to form amyloid fibers *in vitro*. Previously, we found that the fragments of Rnq1_WT_ encompassing the NPD and one or two QN-rich regions of the PD readily formed Thioflavin T (ThT)-binding amyloid fibers with reproducible kinetics and concentration dependence (Kadnar et al., 2010). However, the same assay was much less reproducible for the full-length Rnq1 even though full-length Rnq1 amyloid fibers could be detected by transmission electron microscopy (TEM) similar to fibers formed by shorter Rnq1_WT_ fragments (Vitrenko et al., 2007b; Kadnar et al., 2010).

Since shorter Rnq1 fragments work better for the ThT-binding assay, the ΔB1C2D3_T27P_ construct that carries the Rnq1 NPD, the QG dipeptide repeats, the E helical region and QN4 was made and tested alongside the ΔB1C2D3_WT_ (see Fig 1A for the map of the Rnq1 regions). Both proteins ran according to their expected size on SDS-PAGE gels, although ΔB1C2D3_T27P_ ran slightly higher than ΔB1C2D3_WT_ (Fig 4A). The same change in mobility was observed for proteins carrying the T27P mutations when they were isolated from yeast, indicative of a conformational alteration due to the mutation even in the soluble state (see Fig 2A). Incubation of both proteins in ThT resulted in a shift of the ThT excitation spectrum and increase of ThT fluorescence at 483 nm, indicative of amyloid formation (Fig 4B). Sigmoidal fluorescence kinetics was consistent with the presence of a rate-limiting nucleation step followed by a fiber growth phase for both ΔB1C2D3_T27P_ and ΔB1C2D3_WT_. The length of the lag phase was concentration dependent. Although there was considerable variation between duplicate samples, all showed sigmoidal kinetics and, at equal protein concentrations, the lag phase was similar for the ΔB1C2D3_T27P_ and ΔB1C2D3_wt_ samples.

**Fig 4.**
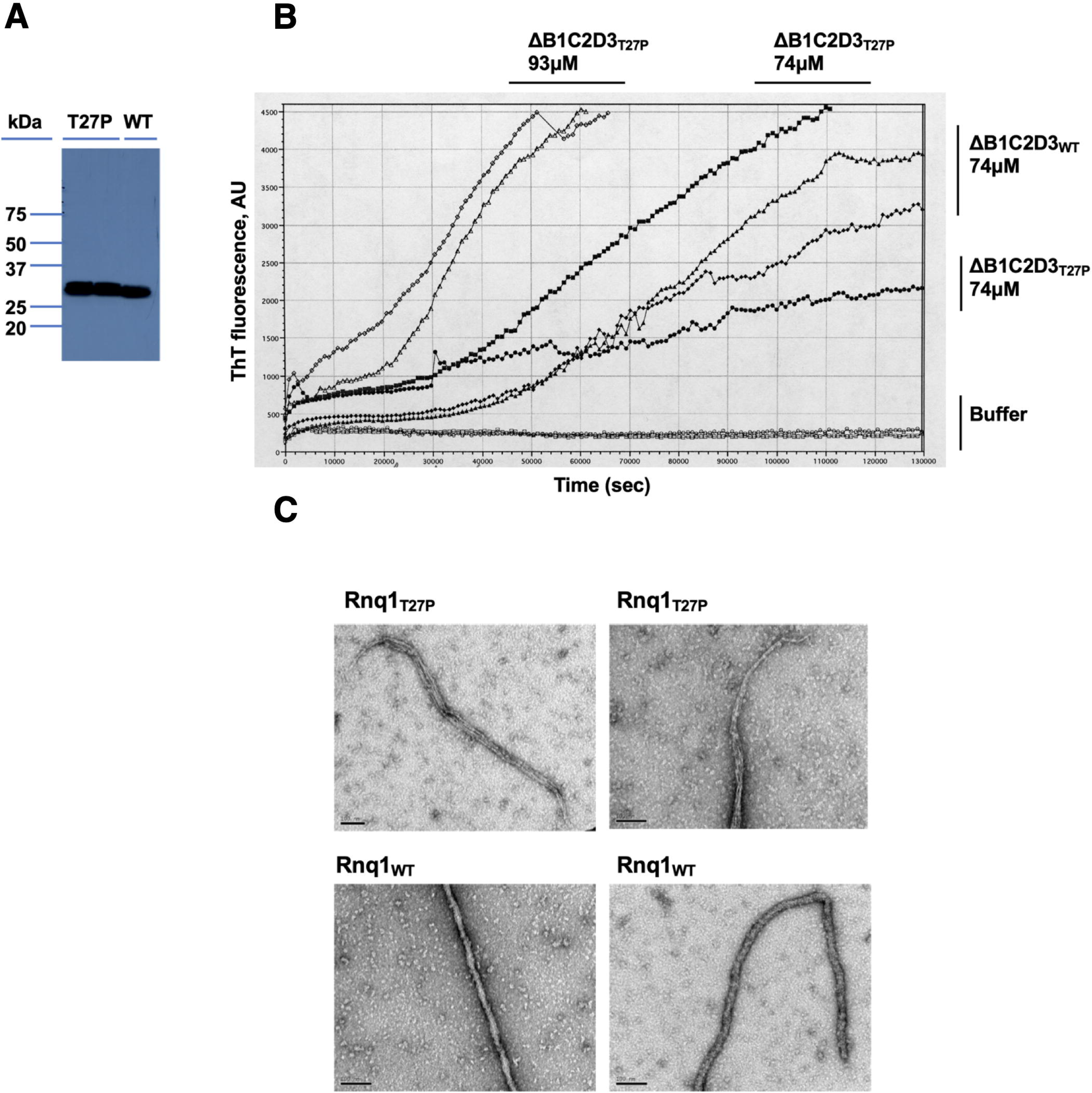
Full-length Rnq1_T27P_ and ΔB1C2D3_T27P_ can form amyloid fibers *in vitro*. **(A)** Western blot analysis of bacterially expressed and purified ΔB1C2D3_T27P_ (T27P; two independent constructs) and ΔB1C2D3_WT_ (WT). The expected size for both constructs is 27kDa. Note slightly slower mobility for ΔB1C2D3_T27P_. Anti-His tag antibody was used for detection. **(B)** Kinetics of *in vitro* aggregation of ΔB1C2D3_T27P_ and ΔB1C2D3_WT_ monitored by ThT fluorescence. Samples and concentrations are indicated on the graph. **(C)** TEM micrographs of negatively stained fibers of Rnq1_T27P_ and Rnq1_WT_. Scale bars are for 100nm. Note lateral association of fibrils in the fibers for Rnq1_T27P_ and twisted appearance of Rnq1_WT_ fibers.

For TEM, full-length Rnq_T27P_ was used and compared to full-length Rnq1_WT_. Fibers were assembled in the presence of ThT for ∼70 hours and an increase of ThT fluorescence was confirmed. TEM revealed >1μm-long non-branched fibers consisting of laterally associated thinner fibrils for both Rnq1_T27P_ and Rnq1_WT_ (Fig 4C). However, the Rnq_T27P_ fibers were less twisted and wider than the Rnq1_WT_ fibers (30 – 50nm *vs* ∼ 20nm) indicating a difference in conformation due to the mutation in the amyloid state.

### The T27P mutation creates a barrier for the transmission of the prion state from [*PIN^+^WT*] to Rnq1T27P

Next, we tested if the T27P mutation affects the transmission of the prion conformation from Rnq1_WT_ to Rnq1_T27P_. If the transmission block were absolute, all post-shuffle cells expressing Rnq1_T27P_ would be [*pin^-^*]. The incomplete Pin^-^ phenotype observed in our experiments is consistent with either the [*PIN^+^_T27P_*] prion being absent in most but not all post-shuffle Rnq1_T27P_ cells, or with the prion state being transmitted and maintained in Rnq1_T27P_ cells but the mutation making [*PIN^+^_T27P_*] extremely weak in its ability to induce [*PSI^+^*]. To distinguish between these possibilities, we changed the protocol from the one where the Pin state of the entire post-shuffle cultures was scored (e.g. in Fig 2B) to a protocol testing the Pin phenotype of individual cells in these cultures (see scheme in Fig 5A). The key difference (circled in red) is that now post-shuffle cultures were colony purified before being tested for the Pin phenotype.

**Fig 5.**
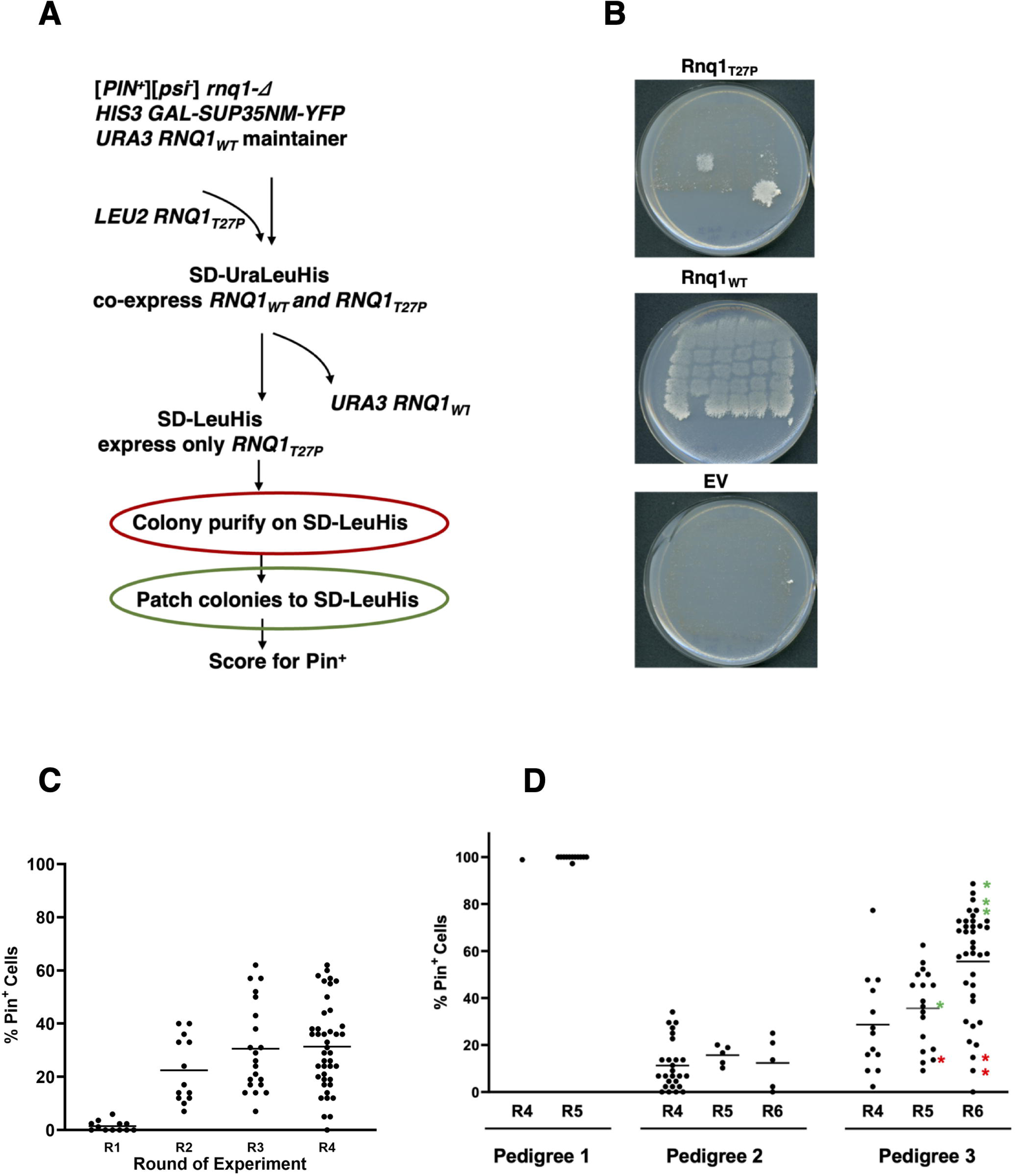
The T27P mutation creates a barrier for the transmission of the prion state from [*PIN^+^_WT_*] to Rnq1_T27P_. (**A**) Scheme of experiments testing the frequency of [*PIN^+^*] loss after substituting Rnq1_T27P_ for Rnq1_WT_ and mitotic stability of [*PIN^+^_T27P_*] isolates. See text. Red circle indicates a step at which yeast were colony purified for the analysis of the fraction of [*PIN^+^_T27P_*] cells in the cultures right after substituting *RNQ1_T27P_* for *RNQ1_WT_* (data shown in Table 2 and illustrated in (B); also corresponds to Round 1 in (C) and (D), and, for other donor and recipient construct combinations, to data in Fig 7B and Round 1 in Figs 7C through 7G). Green circle indicates the step at which [*PIN^+^_T27P_*] isolates were colony purified for the analysis of their mitotic stability (corresponds to data in Table 3 and Round 2 in (C) and (D), and, for other donor and recipient combinations, to Round 2 in Figs 7C through 7G). (**B**) After-shuffle cultures expressing Rnq1_T27P_ contain Pin^+^ and Pin^-^ cells. Sample SD-Ade plates after [*PSI^+^*] induction on SGal-LeuHis with patches expressing the indicated Rnq1 allele or carrying the empty vector. Shown is growth on SD-Ade after 13 days of incubation at 20^0^C. All patches were Ade^-^ after incubation on the control SD-LeuHis media (not shown). **(C)** Increase of mitotic stability of [*PIN^+^_T27P_*] indicative of gradual conformational adaptation of the prion. See text and (A) for the description of the experiment. Data for Round 1 come from colony purification of 12 Rnq1_T27P_ post-shuffle cultures transferred from FOA to SD-LeuHis. Data for Round 2 come from colony purification of the Pin^+^ Rnq1_T27P_ isolates obtained in Round 1; 13 out of 15 obtained isolates were analyzed (the remaining 2 were lost due to contamination). For Round 3, seven Pin^+^ cultures used in Round 2 analysis were randomly chosen, and for each three randomly chosen Pin^+^ isolates were colony purified and scored for Pin^+^ (a total of 21 data points). In Round 4, the analysis was continued for all twelve cultures originating from four out of seven Round 2 cultures (three randomly chosen Pin^+^ isolates for each culture, a total of 36 data points) and for partial lineages of two more Round 2 isolates (a total of 6 more data points). Data for positive (Rnq1_WT_ shuffled to Rnq1_WT_) and negative (Rnq1_WT_ to EV) controls is not included for simplicity, but these experiments were carried through Round 2 on a similar scale and through further rounds on a limited scale. [*PIN^+^_WT_*] maintained ∼100% stability in all isolates tested, with no switching to less stable variants, and yeast with shuffled-in EV did not accumulate Pin^+^ cells. (**D**) Maturation of post-transmission [*PIN^+^_T27P_*] to heritable prion variants with characteristic stabilities. See Results for the description of this experiment and explanation of pedigrees. The R4, R5 and R6 labels correspond to the round of the analysis of mitotic stability of post-shuffle [*PIN^+^_T27P_*]. Asterisks mark directly related parts of the pedigree of the meta-stable [*PIN^+^_T27P_*] prion variant that is prone to yield more stable (red) and less stable (green) [*PIN^+^_T27P_*] variants. Differences between pedigrees 1, 2, and 3 are statistically significant for all rounds of analysis (p<0.01). There are no significant differences within pedigrees 1 and 2 between shown rounds of analysis. For pedigree 3, there is no significant difference between R4 and R5, but for R6 there is a significant difference (p<0.01) due to gradual stabilization of [*PIN^+^_T27P_*] in some isolates (green asterisks).

**Table 2.** Only a small proportion of cells remain Pin^+^ in the *RNQ1_T27P_* expressing cultures after shuffling out the *RNQ1_WT_* plasmid. Shown are results of two independent experiments. For each experiment, colonies from at least 11 post-shuffle Rnq1_T27P_ cultures were tested. T-tests are only shown for Rnq1_T27P_ *vs* EV (for both experiments p<0.05); for all other combinations p<1.0e^-25^. * The only Pin^+^ clone obtained in the EV control in Experiment 2 appears to be a non-Rnq1-based prion.

| Experiment | Construct after plasmid shuffle | Total tested | Pin<sup>+</sup> | Average Pin<sup>+</sup> (%) | SEM (%) | T-test |
| --- | --- | --- | --- | --- | --- | --- |
| # 1 | *RNQ1<sub>WT</sub>* | 435 | 435 | 100 | ± 0 | ] p=0.019 |
|  | *RNQ1<sub>T27P</sub>* | 286 | 7 | 2.23 | ± 0.79 |  |
|  | EV | 197 | 0 | 0 | ± 0 |  |
| # 2 | *RNQ1<sub>WT</sub>* | 420 | 419 | 99.76 | ± 0.23 | ] p=0.038 |
|  | *RNQ1<sub>T27P</sub>* | 796 | 15 | 1.49 | ± 0.53 |  |
|  | EV | 504 | 1* | 0.19 | ± 0.19 |  |

**Table 3.** In the post-shuffle Pin^+^ Rnq1_T27P_ isolates the [*PIN^+^_T27P_*] prion is mitotically unstable and the strength of the Pin^+^ phenotype correlates with fraction of cells retaining the prion. See text and Fig 5A for experimental design (green circle indicates a step at which Pin^+^ isolates were colony purified). The strength of the Pin^+^ phenotype in post-shuffle Rnq1_T27P_ isolates was determined by growth on SD-Ade and SEt-Ade media at 30^0^C and SD-Ade media at 20^0^C after [*PSI^+^*] induction. “Moderate”, “weak” and “very weak” corresponds to the Pin^+^ phenotypes compared to similarly obtained post-shuffle Rnq1_WT_ isolates where the Pin^+^ phenotype is considered “strong” and 100% cells retained the prion. Shown are percentages of Pin^+^ colonies obtained by colony purification of the original Pin^+^ Rnq1_T27P_ isolates, as described in Results.

| Strength of Pin<sup>+</sup> in the original isolate (Round 1) | % Pin<sup>+</sup> cells (Round 2) | Average ± SD |
| --- | --- | --- |
| Very weak | 3.3 | 18.4 ± 12.3 |
| Weak | 14.3 |  |
| Moderate | 27.3 |  |
| Moderate | 28.6 |  |

We found that only 1.5 – 2% of the colonies from the Rnq1_T27P_ post-shuffle cultures remained Pin^+^, whereas the remaining 98 – 98.5% were clearly Pin^-^ (Table 2; also see a sample plate in Fig 5B, but note that this plate was chosen to show the presence of Pin^+^ patches while many plates had no Pin^+^ patches). We additionally confirmed that these Pin^+^ isolates did not retain the *URA3 RNQ1_WT_* plasmid. Also, as expected, the Pin^+^ phenotype depended on the presence of the *LEU2 RNQ1_T27P_* plasmid. In the positive control Rnq1_WT_ post transmission cultures, all but one colony were found to be Pin^+^. This is consistent with the >99.5% [*PIN^+^_WT_*] stability in the original yeast strain before transformation with any plasmids and plasmid shuffle. In the negative control post-transmission cultures carrying just the EV, all but one colony were found to be Pin^-^ (see Table 2 note). The difference between the frequency of Pin^+^ cells in the Rnq1_T27P_ and EV cultures was statistically significant in all experiments (p<0.05). Finally, the 1.5 – 2% Pin^+^ cells detected in the Rnq1_T27P_ post-transmission cultures were not due to *de novo* [*PIN^+^_T27P_*] appearance. This is because [*PIN^+^_T27P_*] does not form in *rnq1-Δ* cells without Rnq1_T27P_ overexpression (see Fig 3B bottom row). Also, when the *LEU2 CEN RNQ1_T27P_* or EV plasmids were introduced into the [*pin^-^*][*psi^-^*] *rnq1-Δ* strain carrying *HIS3 GAL-SUP3NM-YFP*, no Pin^+^ transformants were obtained among ∼1,000 individual transformants tested, and the frequency of Ade^+^ colonies following attempted [*PSI^+^*] induction was no greater in the *RNQ1_T27P_* transformants compared to EV even after several passages on plasmid-selective media.

Taken together, these findings support the idea of a transmission barrier between Rnq1_WT_ and Rnq1_T27P_. If this were true, the proportion of Pin^+^ cells in the initial post-shuffle cultures (Table 2) would be determined by both the frequency of the transmission of the prion state from Rnq1_WT_ to Rnq1_T27P_ and the stability of the resulting [*PIN^+^_T27P_*] prion.

Because the phenotype of the Rnq1_T27P_ Pin^+^ isolates varied in strength (compare the two Rnq1_T27P_ Pin^+^ patches on Fig 5B) and was moderately to significantly reduced relative to Rnq1_WT_ after shorter incubation times or if grown on SD-Ade at 30^0^C (not shown), we hypothesized that the newly forming [*PIN^+^_T27P_*]s were unstable, so the Pin^+^ Rnq1_T27P_ patches contained both [*PIN^+^_T27P_*] and [*pin^-^_T27P_*] cells and the strength of the Pin^+^ phenotype reflected the proportion of the [*PIN^+^_T27P_*] cells. To test this, four Pin^+^ Rnq1_T27P_ isolates were colony purified from the patches made on the SD-LeuHis plasmid-selective media prior to testing for the Pin^+^ phenotype (circled in green in Fig 5A) onto SD-LeuHis, and then individual colonies were patched on the same media and tested for the retention of [*PIN^+^_T27P_*] by [*PSI^+^*] induction. Indeed, in all four Pin^+^ Rnq1_T27P_ isolates both Pin^+^ and Pin^-^ cells were present, indicative of prion instability. Also, as hypothesized, the strength of the Pin^+^ phenotype in the original patch correlated with the fraction of the Pin^+^ cells found (Table 3).

This is reminiscent of what happens when PrP^Sc^ prions cross the inter-species prion transmission barriers (reviewed in Baskakov, 2014) and when Rnq1 fragments encompassing different QN regions of the PD cross intraspecies transmission barriers in yeast (Kadnar et al., 2010). In these cases, crossing the barrier usually leads to poorly propagating prion variants, and stable prion variants are only detected in later passages either after selection and/or conformational adaptation. Thus, we asked if the unstable [*PIN^+^_T27P_*] prions that formed during co-expression of Rnq1_T27P_ with Rnq1_WT_ in [*PIN^+^_WT_*] cells would eventually mature into more stable ones. The stability of [*PIN^+^_T27P_*] was first scored in 13 Pin^+^ Rnq1_T27P_ post-shuffle isolates as described above (Fig 5A, green circle), but this time Pin^+^ isolates were chosen randomly and ∼84 colonies were scored for each isolate to estimate the stability more accurately (Round 2 in Fig 5C). Then, 21 [*PIN^+^_T27P_*] patches obtained in this round were again colony purified from pre-induction SD-LeuHis and patched on SD-LeuHis for Round 3 of [*PIN^+^_T27P_*] stability analysis. This was repeated with 42 Round 3 [*PIN^+^_T27P_*] patches for Round 4 analysis (Fig 5C). The number of cell divisions between each round was equal and estimated to be ∼19. As can be seen, while [*PIN^+^_T27P_*] remained highly unstable in both Round 2 and Round 3 analyses, for some [*PIN^+^_T27P_*] isolates in Round 3 stability increased significantly, with over 60% of cells retaining [*PIN^+^_T27P_*], which was reflected by a moderate increase of the average stability in this round compared to Round 2. In Round 4, however, no dramatic increase in stability was observed, suggesting that, when [*PIN^+^_T27P_*] undergoes maturation without selection, prion variants characterized by different stabilities can be formed and maintained.

To identify such prion variants, we repeated the analysis of post-shuffle [*PIN^+^_T27P_*] stability, this time choosing only three independently obtained [*PIN^+^_T27P_*]s but keeping pedigree data as separate sets and extending analysis beyond Round 4. In Pedigree 1 [*PIN^+^_T27P_*] was extremely unstable early on but yielded an almost 100% stable [*PIN^+^_T27P_*] prion in Round 4, and this prion remained stable in Round 5 (430/431 Pin^+^ in 12 [*PIN^+^_T27P_*] patches tested; Fig 5D). This [*PIN^+^_T27P_*] prion depended on the maintenance of the *RNQ1_T27P_* plasmid, was GuHCl-curable, and promoted the induction of a typical array of [*PSI^+^*] variants at a frequency of a “strong” [*PIN^+^_WT_*]. Such stable [*PIN^+^_T27P_*]s were occasionally detected in other experiments.

In Pedigree 2, [*PIN^+^_T27P_*] remained very unstable in Round 4 and this very low stability was maintained in Rounds 5 and 6, indicative of the establishment of a prion variant not prone to subsequent changes in heritability (Fig 5D). In Pedigree 3 [*PIN^+^_T27P_*] reached a moderate level of stability, significantly higher than in Pedigree 2 (p<0.01; Fig 5D), and remained moderately stable in most cultures in Rounds 5 and 6 but yielded occasional isolates with significantly lower or higher heritable stability (red and green asterisks on Fig 5D), indicative of the establishment of a prion variant that is capable of further conformational adaptation. Altogether, we interpret these data as maturation of post-transmission [*PIN^+^_T27P_*]s to heritable prion variants with characteristic stabilities that is mostly completed within ∼60 cell divisions but may continue longer and go through intermediate variants for some isolates.

### Rnq1T27P can join the amyloid [*PIN^+^*] aggregates formed by Rnq1WT

The strength of transmission barriers is determined by two key factors: the efficiency of transmission of the prion state from the donor prion, [*PIN^+^_WT_*], to the recipient protein, Rnq1_T27P_, and the ability of the recipient protein-based prion, [*PIN^+^_T27P_*], to propagate efficiently. The data presented in the previous section indicate that the inefficient propagation of post-transmission [*PIN^+^_T27P_*]s is definitely a contributing factor to the barrier. After the *RNQ1_WT_* plasmid was shuffled out, [*PIN^+^_T27P_*] was very unstable in all isolates tested and further conformational adaptation was required for [*PIN^+^_T27P_*] stability to increase (see Figs 5C and 5D). We now ask if extreme instability of non-adapted [*PIN^+^_T27P_*] was likely the only reason for presence of only 1.5 - 2% [*PIN^+^_T27P_*] cells in post-shuffle cultures or if other mechanisms were also contributing.

Here, we tested if Rnq1_T27P_ could join the pre-existing [*PIN^+^_WT_*] prion aggregates because a block at this step would inhibit the transmission of the prion state regardless of the resulting stability of [*PIN^+^_T27P_*] prions. The Rnq1_WT_-CFP and Rnq1_T27P_-YFP were co-expressed in a [*PIN^+^_WT_*] strain from *CEN* plasmids where constructs were controlled by the *CUP1* promoter. To make sure that no significant *de novo* induction of [*PIN^+^_T27P_*] has occurred before cell analysis, cultures of co-transformants were grown to mid-log in the plasmid-selective no-copper media, and CuSO_4_ was added to 5μM only 2 hours prior to microscopy (see Materials and Methods). At this low concentration of CuSO_4_, Rnq1 is expressed at levels approximately equal to those of chromosomally encoded Rnq1 (Fig S3). As shown in Fig 6A, Rnq1_WT_-CFP and Rnq1_T27P_-YFP co-localized in bright fluorescent foci in most cells, indicative of Rnq1_T27P_ being able to join [*PIN^+^_WT_*] aggregates. We confirmed that Rnq1_T27P_-YFP joins [*PIN^+^*] aggregates just as efficiently as Rnq1_WT_-CFP (Figs S2A and S2B). Co-localization was retained even after prolonged incubation or a higher-level expression of the constructs (5μM CuSO_4_ overnight, 20μMcuSO_4_ for 2 hours or overnight, or 50μMcuSO_4_ for 4 hours; not shown). The appearance of Rnq1_T27P_-YFP foci was indistinguishable from that of Rnq1_WT_-CFP foci and corresponded to a foci pattern characteristic of the “multi-dot high” [*PIN^+^*] variant (Bradley and Liebman, 2003) used for these experiments. Also, there was no evidence suggesting that Rnq1_T27P_-YFP aggregates were just seeded by Rnq1_WT_ but then formed next to pre-existing [*PIN^+^*] aggregates not incorporating Rnq1_WT_: the overlap of Rnq1_T27P_-YFP and Rnq1_WT_-CFP foci was usually complete in the focal plane imaged (Figs 6A, S2A, and S2B) and in the focal planes immediately above and below. The foci with co-localized Rnq1_WT_-CFP and Rnq1_T27P_-YFP were not sensitive to 1,6 hexanediol (Hex; Fig 6A), with Rnq1_T27P_ retaininged after the treatment, suggesting that the fluorescently tagged proteins were joining the typical amyloid [*PIN^+^_WT_*] aggregates (see also Fig 8A for ThT staining). Thus, the transmission barrier is not due to the inability of Rnq1_T27P_ to join pre-existing [*PIN^+^_WT_*] aggregates formed by Rnq1_WT_ or to convert into amyloid.

**Fig 6.**
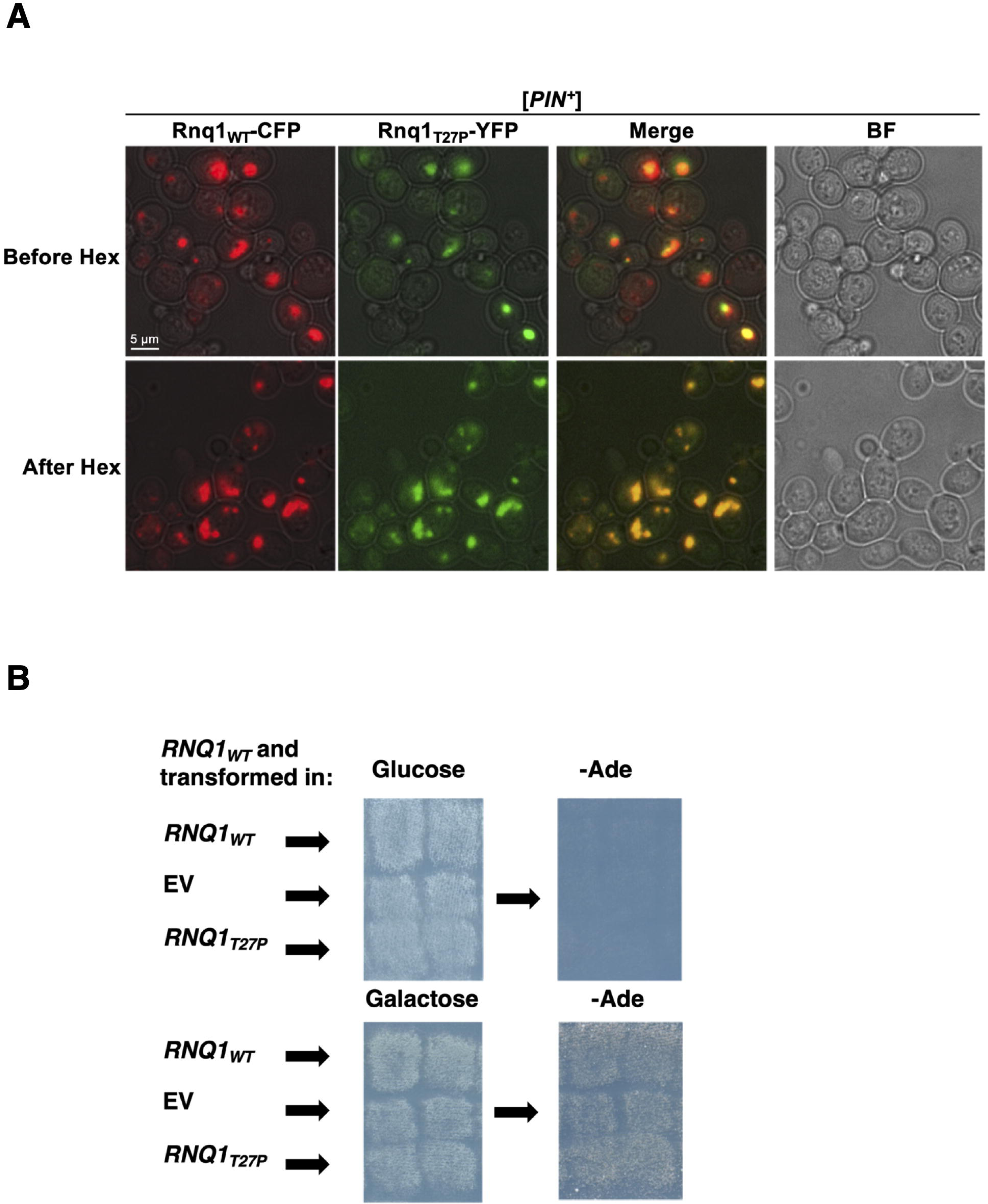
The transmission barrier between [*PIN^+^_WT_*] and Rnq1_T27P_ is not caused by the inability of Rnq1_T27P_ to co-aggregate with Rnq1_WT_ or by loss of the [*PIN^+^*] prion during co-expression of Rnq1_WT_ and Rnq1_T27P_. (**A**) Rnq1_T27P_ can join the [*PIN^+^_WT_*] prion aggregates. The [*PIN^+^*] 74-D694 strain was co-transformed with the *CEN LEU2 CUP1-RNQ1_WT_-CFP* and *CEN URA3 CUP-RNQ1_T27P_-YFP* plasmids. Cultures of co-transformants were grown overnight (∼to mid-log) in no-copper SD-LeuHis media, diluted to OD_595_=1.0 into SD-LeuHis+5μM CuSO_4_ and grown for 2 more hours prior to observing the cells with or without Hex treatment. A representative group of cells is shown. See Fig S2A for an image of a larger field and analysis of Rnq1_WT_ and Rnq1_T27P_ co-localization individual cells. (**B**) Rnq1_T27P_ does not affect the Pin^+^ phenotype when co-expressed with Rnq1_WT_ in a [*PIN^+^*] stain. Shown are the images of the transformants of the YID146.3 cells carrying the *GAL-SUP35NM-YFP* [*PSI^+^*] inducing construct along with the indicated plasmid combinations. “Galactose” and “Glucose” indicate SGal-UraLeuHis and SD-UraLeuHis media, respectively. Images on these media were taken after 48 hours of growth at 30^0^C. From these media yeast were replica plated to SD-Ade. Images were taken after 14 days at 30^0^C. Note that these are patches from the plates shown on Fig 2B and Fig S1 but before plasmid shuffle.

**Fig 7.**
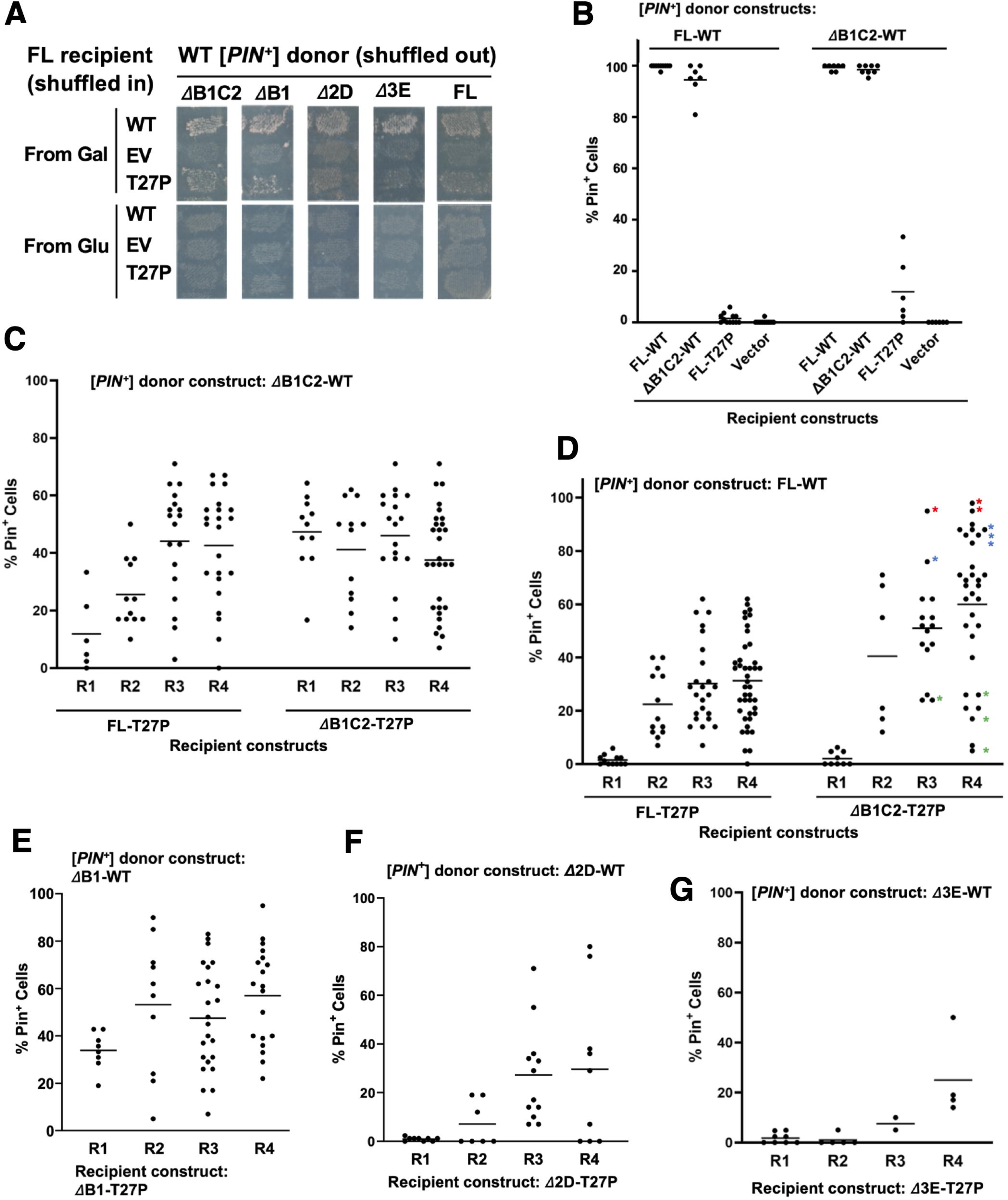
The transmission barrier created by the T27P mutation in the NPD is imposed through the first two QN-rich regions in Rnq1 PD. **(A)** Deletion of B1 and B1C2 but not 2D and 3E regions in the PD of Rnq1_WT_ reduces the transmission barrier from [*mini-PIN^+^_WT_*]s towards Rnq1_FL-T27P_. The [*PSI^+^*] induction test was done in cultures where the Rnq1_FL-WT_, Rnq1_FL-T27P_ recipient constructs (or EV) were substituted for the indicated deletion constructs in the [*mini-PIN^+^_WT_*] or [*PIN^+^_FL-WT_*] prion donor strains. Shown are images of growth on SD-Ade plates replica plated from either the [*PSI^+^*] inducing SGal-LeuHis or the control glucose media and grown for 10 days at 30^0^C. **(B – G)** Analysis of the role of QN-rich regions in the NPD of Rnq1 on the transmission barriers and subsequent conformational adaptation of Rnq1-based prions in the presence of the T27P mutation in the NPD. Experiments were performed as described in Fig 5. Donor and recipient constructs for prion transmission are indicated; nomenclature marks the deleted regions within the PD (ΔB1C2, ΔB1, Δ2D, or Δ3E; see Fig 1A) compared to the full-length Rnq1 protein (FL) and the presence or absence of the T27P mutation in the construct (T27P or WT, respectively). Shown is the fraction of prion-carrying cells in the cultures after plasmid shuffle (B and R1 in C – G) and mitotic stability of the resulting prions during further propagation of randomly chosen isolates (R2 through R4; ∼19 cell generations between each round of analysis). In (D) asteriscs indicate related progeny for cultures in R3 and R4 highlighting the establishment of [*PIN^+^_τιB1C2_*] variants with high (red and blue) and low (green) mitotic stability. **Experiments in (B – G) were conducted in parallel, so all data can be compared between the graphs.** Some control experiments (e.g. analysis of transmissions between non-mutant constructs and lack of transmissions in substitutions for EV) are omitted on graphs for simplicity: the results were as expected and in full agreement with Kadnar et al. (2010). All conclusions in Results are based on statistically significant differences or lack of thereof (0.05 or 0.01 probability cutoff).

**Fig 8.**
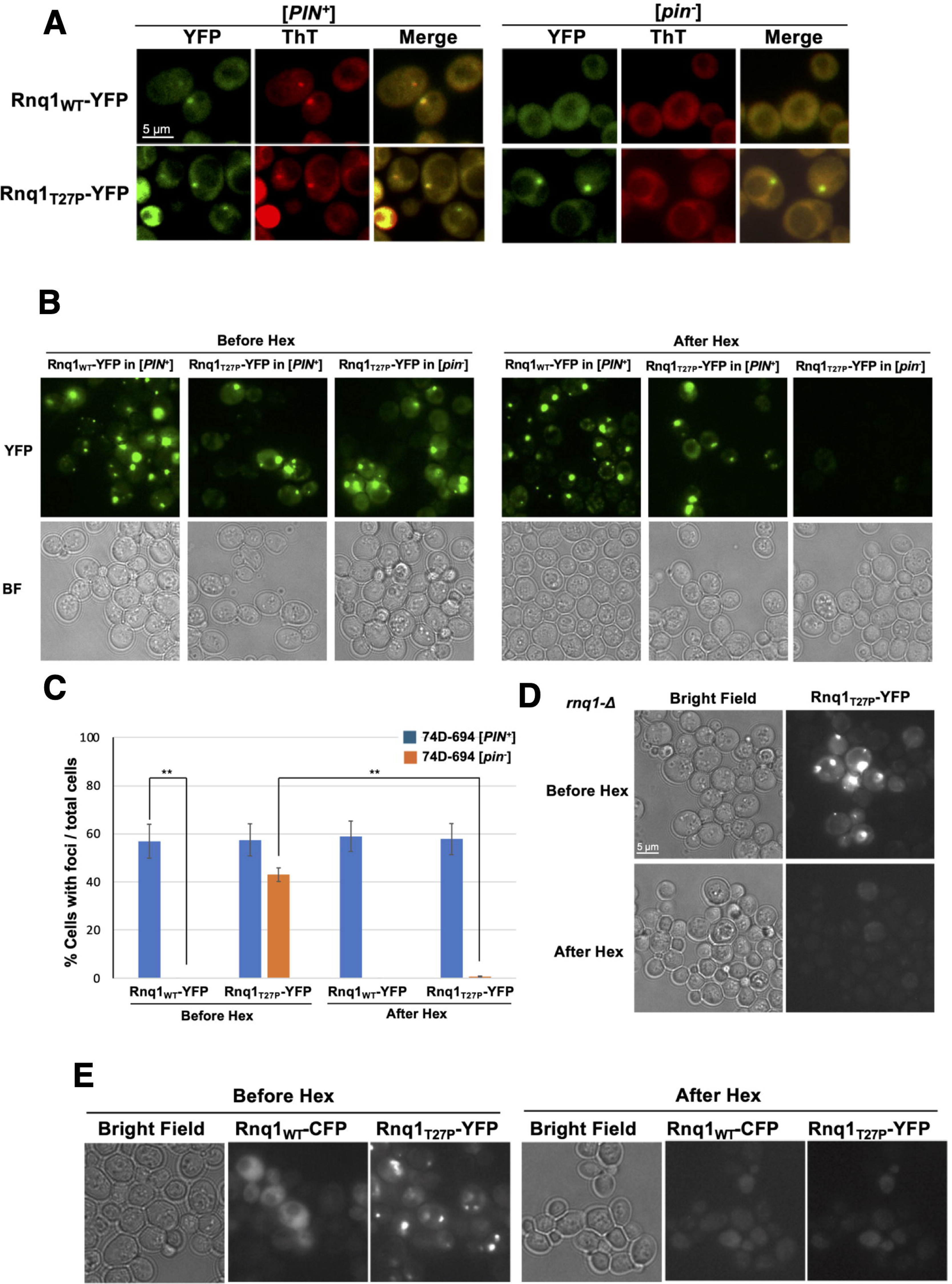
The T27P mutation in the NPD of Rnq1 promotes the formation of liquid-like droplets by Rnq1_FL-T27P_-YFP expressed in [*pin^-^*] cells even at levels close to the expression of endogenous Rnq1. (A) Unlike Rnq1_FL-WT_-YFP, Rnq1_FL-T27P_-YFP forms aggregates in [*pin^-^*] cells and these aggregates are not amyloid. Cultures of isogenic [*PIN^+^*] and [*pin^-^*] strains transformed with plasmids carrying the indicated constructs under the control of *CUP1* promoter were grown overninght in SD plasmid selective media to OD_595_ ∼ 0.8. Then expression of the constructs was induced by addition of CuSO_4_ to 50μM and growth was continued for 4 more hours prior to ThT staining and imaging. **(B - C)** Aggregates formed by Rnq1_FL-T27P_-YFP in [*pin^-^*] cells are liquid-like droplets. Yeast transformants were grown overnight in liquid SD media selective for plasmids. Then OD_595_ was adjusted to 1.0, CuSO_4_ was added to 5μM, and cultures were grown for two more hrs before Hex treatment. (C) is quantification of data in experiment illustrated in (B). Asterisks indicate statistically significant differences (p<0.01). **(D)** Rnq1_FL-T27P_-YFP forms liquid-like droplets in *rnq1-Δ* cells. Experiment was performed as described for (B - C). **(E)** Rnq1_FL-WT_-CFP does not join liquid-like droplets formed by Rnq1_FL-T27P_-YFP in [*pin^-^*] cells. The [*pin^-^*] strain was co-transformed with the *CEN LEU2 CUP1-RNQ1_WT_-CFP* and *CEN URA3 CUP-RNQ1_T27P_-YFP* plasmids. Experiment was performed exactly as in Fig 6A where Rnq1_FL-T27P_-YFP is shown co-aggregate with Rnq1_FL-WT_-CFP in [*PIN^+^_WT_*] cells.

### The co-expression of Rnq1T27P does not lead to loss of [*PIN^+^WT*]

We then asked if introduction of the Rnq1_T27P_ expressing plasmid led to a rapid loss of [*PIN^+^_WT_*] prior to the plasmid shuffle. One possibility is that incorporation of Rnq1_T27P_ into the [*PIN^+^_WT_*] aggregates “poisoned” these aggregates and blocked the prion propagation. This hypothesis was used to explain the curing of the [*URE3*] prion upon the overexpression of Ure2-GFP or some Ure2 fragments (Edskes et al., 1999) and curing of [*PSI^+^*] by dominant “[*PSI^+^*] no more” *PNM* mutations within Sup35 PD (Kochneva-Pervukhova et al., 1998; Derkatch et al., 1999; DiSalvo et al., 2011; Pei et al., 2017).

However, we have found that transforming the [*PIN^+^_WT_*] strain used in the genetic screen ([*PIN^+^*] [*psi^-^*] *rnq1-Δ* carrying the *CEN URA3 RNQ1_WT_* [*PIN^+^*] maintainer and the *CEN HIS3 GAL-SUP35NM-YFP* [*PSI^+^*] inducer; see Fig 1B) with the *CEN LEU2 RNQ1_T27P_* plasmid did not lead to loss of the [*PIN^+^*] prion. All 240 transformants tested in this “no shuffle” experiment remained Pin^+^ even after four more passages on media selective for all the plasmids. Furthermore, no reduction of [*PSI^+^*] induction was observed in such cells compared to the control transformants with *CEN LEU2 RNQ1_WT_* or *CEN LEU2* EV in the nonsense suppression test (Fig 6B) and when *de novo* induction of [*PSI^+^*] was measured as a fraction of cells with Sup35NM-YFP aggregates during overexpression of the Sup35NM-YFP construct (regardless of the plasmid introduced, ∼50% cells had line-, ring-, and dot-shaped aggregates after 48 hours of growth on SGal-UraLeuHis media; data not shown).

Also, there was no indication of incompatibility of Rnq1_WT_ and Rnq1_T27P_: there was no growth inhibition for the *RNQ1_T27P_* carrying cells compared to the *RNQ1_WT_* and EV transformants (Fig 6B). The stability of the [*PIN^+^_WT_*] maintainer plasmid, *CEN URA3 RNQ1_WT_*, was the same in the *CEN LEU2 RNQ1_T27P_*, *CEN LEU2 RNQ1_WT_*, and *CEN LEU2* EV transformants (AVE ± SD are 38.4 ± 11.3%, 41.9 ± 10.3%, and 41.6 ± 5.8%, respectively). Also, there was no difference in the stability of the *CEN LEU2 RNQ1_T27P_*, *CEN LEU2 RNQ1_WT_*, and *CEN LEU2* EV (AVE ± SD are 39.9 ± 7.8%, 34.8 ± 10.2%, and 38.0 ± 7.7%, respectively; see Materials and Methods for the description of the test).

We also analyzed if similar or different arrays of [*PSI^+^*] variants were induced by Sup35NM-YFP overexpression in cells co-expressing Rnq1_WT_ and Rnq1_T27P_ compared to cells expressing two copies of Rnq1_WT_. For either the cells co-expressing Rnq1_WT_ and Rnq1_T27P_ or cells expressing two copies of Rnq1_WT_ most [*PSI^+^*] variants were moderate (76.1 and 89.7%, respectively), and strong [*PSI^+^*] variants were rare (∼3%). However, the percentage of weak [*PSI^+^*] variants was significantly higher in cells co-expressing Rnq1_T27P_ and Rnq1_WT_ (21.0 *vs* 6.5%; p <0.01; Table 4).

**Table 4.** Co-expression of Rnq1_T27P_ and Rnq1_WT_ leads to a slight shift towards the induction of weak [*PSI^+^*] variants compared to expression of two copies of Rnq1_WT_. Following the induction of [*PSI^+^*] by high level expression of Sup35NM-YFP on SGal-UraLeuHis media selective for all plasmids, yeast were replica plated on glucose SD-Ade media not selective for any plasmid and incubated at 20^0^C for 25 days. Then Ade^+^ cells were scraped from the SD-Ade plates and colony purified on YPD. Random individual colonies were patched on YPD before color developed. [*PSI^+^*]s isolated from at least four independent transformants carrying the indicated plasmids were analyzed by the color test on YPD, growth on adenineless media and curability by GuHCl. Results of all three tests were consistent for ∼99% [*PSI^+^*] variants.

| Plasmid introduced | [*PSI*<sup>+</sup>] variants: number isolated (% ± SD) |  |  |  |
| --- | --- | --- | --- | --- |
|  | Weak | Medium | Strong | Total |
| *RNQ1<sub>WT</sub>* | 9 (6.5 ± 5.5) | 125 (89.7 ± 8.9) | 5 (3.8 ± 6.6) | 139 (100) |
| *RNQ1<sub>T27P</sub>* | 15 (21.1 ± 5.1) | 54 (76.1 ± 4.9) | 2 (2.8 ± 3.2) | 71 (100) |
| EV | 2 (3.1 ± 3.6) | 64 (96.9 ± 3.6) | 0 (0 ± 0) | 66 (100) |

Thus, the barrier for the transmission of the [*PIN^+^_WT_*] prion state to Rnq1_T27P_ is not due to Rnq1_T27P_ being incompatible with Rnq1_WT_ co-expression or “poisoning” of the [*PIN^+^_WT_*] prion aggregates, but upon joining [*PIN^+^_WT_*] aggregates Rnq1_T27P_ changes the ability of the Rnq1-based prion to promote the induction of particular [*PSI^+^*] variants.

### The transmission barrier created by the T27P mutation in the NPD is eliminated by deletion of the first two QN-rich regions in Rnq1 PD

Data presented in the previous sections indicate that the major reason for the transmission barrier is the inability of Rnq1_T27P_ to attain an efficiently propagating prion conformation when seeded by the [*PIN^+^_WT_*] variant used in our studies. We now ask if other conformational variants of Rnq1-based prions would be more acceptable donors of the prion state for Rnq1_T27P_. In our earlier studies we uncovered transmission barriers with subsequent conformational adaptation for the transfer of prion conformations between the [*PIN^+^_WT_*] variant used in the current study and Rnq1 fragments lacking one or more QN-rich regions in the PD (Kadnar et al., 2010; see Fig 1A). The stable prion variants that were formed by these Rnq1 fragments after crossing the transmission barrier from full-length Rnq1 (Rnq1_FL_) and consequent conformational adaptation were collectively named [*mini-PIN^+^*]s and were confirmed to have prion conformations different from those of [*PIN^+^_FL_*]. Thus, we tested if the transmission barrier to Rnq1_T27P_ was reduced when these stable [*mini-PIN^+^*]s not carrying the T27P mutation were used as donors of the prion conformation instead of full length [*PIN^+^_WT_*]. From now on, the **nomenclature** will include both the presence of full-length Rnq1 or of a Rnq1 fragment, as well as the WT or T27P allele (e.g. [*PIN^+^_FL-WT_*] or [*PIN^+^_FL-T27P_*]). For the Rnq1 fragments, the nomenclature reflects the deleted part of the PD, which includes the QN-rich region(s) numbered as in Fig 1A and an adjacent helical stretch(es) indicated in Fig 1A by letters (e.g. [*PIN^+^_τιB1C2-WT_*] or [*PIN^+^_τιB1C2-T27P_*]).

The plasmid shuffle experiment described in Fig 1B was repeated using stable [*mini-PIN^+^_WT_*] stains instead of [*PIN^+^_FL-WT_*]. According to the nonsense suppression-based [*PSI^+^*] induction test, transmission of the prion state to Rnq1_FL-T27P_ from [*PIN^+^_τιB1C2-WT_*] and [*PIN^+^_τιB1-WT_*] but not from [*PIN^+^_τι2D-WT_*] or [*PIN^+^_τι3E-WT_*] was considerably more efficient compared to transmission from [*PIN^+^_FL-WT_*] (Fig 7A). In the case of transmission from [*PIN^+^_τιB1C2-WT_*], a colony purification experiment done as described in Fig 5A was used to confirm that growth on adenineless media in Fig 7A was indeed due to a higher fraction of [*PIN^+^_FL-T27P_*] cells in the “post-shuffle” cultures (Fig 7B; the difference is statistically significant, p<0.01). This supports the hypothesis that the T27P mutation in the NPD changes the range of prion conformations permissible for PD and that [*PIN^+^_τιB1C2-WT_*] and [*PIN^+^_τιB1-WT_*] fall into this range of acceptable conformations while [*PIN^+^_FL-WT_*] does not.

Considering the indication that deletion of the B1C2 region of the Rnq1 PD reduced the transmission barrier, we asked if this region fully determined the inability of Rnq1_FL-T27P_ to take on the prion conformation compatible with [*PIN^+^_FL-WT_*]. The T27P mutation was introduced into the Rnq1 fragment lacking the B1C2 region (the Rnq1_τιB1C2-T27P_ construct), and transmission of the prion state to this construct was assessed using [*PIN^+^_τιB1C2-WT_*] as a donor of the prion state. We found that a significantly higher fraction of Pin^+^ cells was present in the “post-transmission” cultures when both the donor and recipient of the prion state lacked the B1C2 region (Fig 7C; p<0.01). However, no subsequent change of mitotic stability was observed following such transmissions: after four rounds of consecutive colony purifications done as in Fig 5C, the [*PIN^+^_τιB1C2-T27P_*] isolates retained the same stability as immediately after the transmission (Fig 7C, no statistically significant difference between R1 and R2 through R4). Furthermore, one more round of analysis of R4 [*PIN^+^_τιB1C2-T27P_*] isolates with the highest fraction of prion-retaining cells did not indicate that these isolates underwent a heritable conformational shift leading to higher mitotic stability (not shown). One explanation for these data is that the T27P mutation limits the fragmentation of all or most [*PIN^+^_τιB1C2-T27P_*] prion variants thus preventing them from propagating efficiently enough to achieve high mitotic stability. The alternative explanation is that there is no transmission barrier between [*PIN^+^_τιB1C2-WT_*] and Rnq1_τιB1C2-T27P_. In this case, transmission of the prion state from [*PIN^+^_τιB1C2-WT_*] to Rnq1_τιB1C2-T27P_ will result in all cells attaining the same heritable [*PIN^+^_τιB1C2-WT_*] prion conformation with a particular mitotic stability (that happens to be ∼45% after ∼19 cell generations), and then this [*PIN^+^_τιB1C2-T27P_*] variant propagates faithfully without undergoing any further conformational adaptation. Because there is no selection favoring the maintenance of the [*PIN^+^_τιB1C2-T27P_*] prion in our experimental system, we do not expect to detect rare switches to mitotically stable variants within the relatively small sample sizes tested. Also, the proportion of Pin^+^ cells in Round 1 analysis is expected to reflect the 100% efficient transmission of the prion state between [*PIN^+^_τιB1C2-WT_*] and Rnq1_τιB1C2-T27P_ and then the loss of [*PIN^+^_τιB1C2-T27P_*] in cells divisions between the elimination of the *RNQ1_τιB1C2-WT_* plasmid and the start of the analysis of “post-shuffle” cultures, which is ∼18-20 cells divisions, i.e. similar to the ∼19 number of divisions between subsequent rounds.

To distinguish between these two explanations, transmission of the prion state to Rnq1*_τι_*_B1C2-T27P_ was attempted using [*PIN^+^_FL-WT_*] as a donor. Data in Fig 7D indicate the existence of a strong transmission barrier, similar to that between [*PIN^+^_FL-WT_*] and Rnq1_FL-T27P_, and subsequent conformational adaptation which allowed for the establishment of [*PIN^+^_τιB1C2-T27P_*] variants with different levels of mitotic stability, including high mitotic stability that was maintained in subsequent rounds of analysis (see asterisks in Fig 7D). This excludes the explanation that the T27P mutation makes [*PIN^+^_τιB1C2-T27P_*] variants intrinsically unstable and supports the alternative explanation that the T27P mutation in the NPD creates a transmission barrier through the effect on the conformational flexibility of the B1C2 region of the PD. Comparison of data presented in Figs 7C and 7D supports the idea that when both the donor and the recipient lack this region, the mutant attains the prion conformation without a barrier but the mutation slightly inhibits the stability of the resulting [*PIN^+^_τιB1C2-T27P_*] variant. The deletion of the B1C2 region only in the T27P carrying recipient and not in the non-mutant donor is not sufficient to eliminate the transmission barrier. However, in transmissions from [*PIN^+^_FL-WT_*], the subsequent increase of mitotic stability was significantly faster for [*PIN^+^_τιB1C2-T27P_*] compared to [*PIN^+^_FL-T27P_*] (Fig 7D), indicative of the effect of the B1C2 region of PD on the conformational adaptation properties of T27P mutants.

Because the [*PSI^+^*] induction test in Fig 7A indicated that deletion of only the B1 region within the PD could also reduce the transmission barrier, we asked if the first QN-rich region of the PD, QN1, was solely responsible for the effect on both the WT to T27P transmission barrier and conformational adaptation of the T27P carrying prions observed for ΔB1C2 fragments. This is particularly interesting because QN1 is very short, only 12aa (NSNNNNQQGQNQ). Thus, the T27P mutation was introduced into the Rnq1 fragment lacking QN1 but retaining QN2 (the Rnq1_τιB1-T27P_ construct) and the [*PIN^+^_τιB1-WT_*] donor was used to transmit the prion state to this construct. Like in the similar experiments with WT and T27P carrying ΔB1C2 Rnq1 fragments, the fraction of [*PIN^+^_τιB1-T27P_*] cells in the “post-shuffle” cultures was significantly increased compared to the transmissions between FL constructs (p<0.01; compare Fig 7E with 7D), even though the fraction of Pin^+^ cells was slightly lower in [*PIN^+^_τιB1-T27P_*] than in [*PIN^+^_τιB1C2-T27P_*] samples (p<0.05; compare Figs 7C and 7E). However, in the [*PIN^+^_τιB1-T27P_*] isolates, subsequent appearance of [*PIN^+^_τιB1-T27P_*]s with high mitotic stability, indicative of the ongoing conformational adaptation, led us to conclude that removal of only the B1 region in both the donor and the recipient did not eliminate the transmission barrier created by the T27P mutation. This finding suggested that the QN2 region of the Rnq1 PD was also involved in the interaction of the PD with the T27P mutation in the NPD. Yet, similar experiments attempting to transmit the prion state from the [*PIN^+^_τι2D-T27P_*] donor to the Rnq1_τι2D-T27P_ recipient did not suggest that deleting the QN2 region reduced the transmission barrier or led to the establishment of the T27P carrying prions not prone to further conformational adaptation (Fig 7F). Thus, the NPD exerts its effect on the QN1 and QN2 regions of the PD acting in concert rather than individually. Finally, consistent with data shown in Fig 7A, we obtained no evidence suggesting that the deletion of the QN3 region of the Rnq1 PD (Δ3E) could remove the transmission barrier created by the T27P mutation or affect conformational adaptation of [*PIN^+^_τι_*_3E-T27P_] prion variants after crossing the barrier (Fig 7G).

### Unlike Rnq1WT-YFP, Rnq1T27P-YFP readily forms liquid-like droplets when expressed in [*pin^-^*] cells

While Rnq1 aggregation in [*PIN^+^*] strains can be visualized by expression of Rnq1_WT_, its PD, or PD fragments fused to C-terminal fluorescent tags from the GFP family (Sondheimer and Lindquist, 2000; Derkatch et al., 2001; Osherovich and Weissman, 2001; Vitrenko et al., 2007b; Kadnar et al., 2010), overexpression of Rnq1_WT_ fused to such tags (e.g. Rnq1_WT_-GFP) in [*pin^-^*] strains does not lead to the formation of visible aggregates and to the *de novo* appearance of [*PIN^+^*] (Derkatch et al., 2001; Kadnar et al., 2010). One possible explanation is that the extreme C-terminus of the Rnq1 PD encompassing the last QN-rich region, QN4 (see Fig 1A), is important during the step of the *de novo* [*PIN^+^*] prion formation, and bulky C-terminal tags interfere with this process (Galliamov et al., 2024). However, once [*PIN^+^*] is formed, other QN-rich regions within the RNQ1 PD get engaged in the maintenance of the prion structure and blocking of QN4 by a tag is not critical for the prion maintenance (Kadnar et al., 2010).

Because our data indicate that the T27P mutation changes the conformational flexibility of the Rnq1 PD through particular QN-rich regions, we asked if, in the presence of T27P, the C-terminally tagged constructs retain aggregation properties despite the tag partially blocking the QN4 region. As seen in Fig 8A, in contrast to Rnq1_WT_, when Rnq1_T27P_-YFP is expressed in [*pin^-^*] cells, it readily forms dot aggregates, one or several dots per cell. Such dot-like aggregates were formed in ∼40% of cells even when Rnq1_T27P_-YFP was expressed briefly and at close to physiological levels of expression (5μM CuSO_4_ for 2 hours; Figs 8B and 8C; also see Fig S3 for protein expression).

These dot aggregates are clearly distinct from ribbon-like aggregates of the *de novo* forming [*PIN^+^_WT_*]s when GFP-tagged Rnq1 is overexpressed in [*URE3*] cells (Derkatch et al., 2001). Therefore, we tested if Rnq1_T27P_-YFP aggregates forming in [*pin^-^*] cells are amyloid, and found they are not because the aggregates are not stained by Thioflavin T (Fig 8A). Furthermore, these aggregates are dissolved by a short-term Hexanediol (Hex) treatment, suggesting that they are liquid-like droplets (Figs 8B and 8C). The presence of Rnq1_WT_ is not required for Rnq1_T27P_ to form liquid-like droplets as they also form in *rnq1-Δ* cells (Fig 8D). Finally, Rnq1_WT_-CFP does not join the Rnq1_T27P_-YFP when co-expressed in [*pin*^-^] cells (Fig 8E). This further supports the idea that the NPD mutation T27P modulates aggregation and conformational properties of QN-rich regions within the PD of Rnq1.

## Discussion

### The T27P mutation in the NPD of Rnq1 creates a barrier for the transmission of the [*PIN^+^*] prion state from Rnq1WT

Most known prion-forming proteins have chimeric organization containing: 1) a functional non-prion domain, NPD, that is not essential for prion formation and is required for the protein to perform its main cellular function; 2) a prion domain, PD, which is sufficient for prionization and can form prions in the absence of NPD and even when attached to other proteins. At first glance, Rnq1 shares such chimeric organization except that its NPD is very short, compared to its PD, which constitutes up to 2/3 of the protein and has a very complex organization (see Introduction and Fig 1A). The length of the NPD is ∼132 – 153 aa but is not known exactly because *S. cerevisiae* Rnq1, as well as its potential homologs from other species have no known function and are poorly conserved (ILD unpublished observations).

We aimed at understanding the structure - function of Rnq1 related to its prion-forming abilities. Two approaches were used. Previously, we performed extensive deletion analysis of all potential structural features within the PD (Kadnar et al., 2010). Here, we used random mutagenesis of *RNQ1*. We mutagenized both the PD and NPD because, first, the exact length of the NPD is not known and, second, several lines of evidence indicated that the NPD is involved in [*PIN^+^*] formation and maintenance (see below). The genetic screen was for the loss of the Pin^+^ phenotype, *i.e.* the ability of [*PIN^+^*] to facilitate the *de novo* induction of [*PSI^+^*]. This readout allowed screening for mutations in the context of the complete Rnq1 not fused to reporters that were previously shown to modify its prion forming ability (Sondheimer and Lindquist, 2000; Derkatch et al., 2001; Bardill and True, 2009). Here we describe the Rnq1_T27P_ mutant obtained in this screen. We propose that, through interactions with specific QN-rich regions in the PD, this NPD mutation alters the array of conformations the prion domain can attain and thus creates a transmission barrier with a [*PIN^+^_WT_*] prion formed by non-mutant Rnq1. The mutation also alters the aggregation properties of Rnq1 by promoting the formation of liquid-like droplets when Rnq1_T27P_ is fused to a C-terminal YFP tag.

The T27P mutation led to the loss of the [*PIN^+^*] prion in 98 – 98.5% of the cells when Rnq1_T27P_ was substituted for Rnq1_WT_ in a [*PIN^+^_WT_*] strain (Table 2). We concluded that this occurs because the mutation creates a barrier for the transmission of the prion state from Rnq1_WT_ to Rnq1_T27P_ after not finding support for the alternative explanation that Rnq1_T27P_ is intrinsically defective in prion formation and propagation. Indeed, Rnq1_T27P_ can form amyloid *in vitro* (Fig 4C). Also, transient overexpression of Rnq1_T27P_ in a *rnq1-Δ* strain leads to a heritable Pin^+^ phenotype indicative of the *de novo* formation of [*PIN^+^_T27P_*] (Fig 3). Finally, a [*PIN^+^_T27P_*] variant with mitotic stability of >99.9% was obtained upon prolonged mitotic propagation of an unstable [*PIN^+^_T27P_*] retrieved immediately after transmission of the prion state from [*PIN^+^_WT_*] (Fig 5D).

The concept of transmission barriers dates to observations that experimental transmission of scrapie between different mammalian species is frequently far less efficient than within the same species (Pattison et al., 1965). These barriers were then attributed to differences in sequences of the PrP protein in different species (Scott et al., 1989). Interspecies transmission barriers were also uncovered for [*PSI^+^*] and [*URE3*] (Santoso et al., 2000; Chen et al., 2007, 2010; Edskes et al., 2009; Afanasieva et al., 2011; Sharma et al., 2016; Wickner et al., 2019). Importantly, the analysis of these barriers focused on the conformational constraints resulting from sequence differences **exclusively within the PDs** of these proteins. In closely related *Saccharomyces sensu stricto* species, transmission barriers formed even though PDs are highly homologous and carry only a few individual amino acid changes (Santoso et al., 2000; Chen et al., 2007, 2010). Furthermore, mutagenesis of Sup35 and Rnq1 and analysis of Sup35 from wild isolates revealed that transmission barriers can be created within *S. cerevisiae* by single mutations in the PDs (DePace et al., 1998; Bardill and True, 2009; Bateman and Wickner 2012).

However, differences in amino acid sequences in the prion-forming regions are not the only contributing factor to transmission barriers. Studies of mammalian prions eventually concluded that the efficiency of interspecies transmissions is determined by both the PrP sequence and particular PrP^Sc^ conformations in the donor species (Hill et al., 1997; Hill and Collinge 2001; Collinge and Clarke, 2007; Collinge, 2012). Our analysis of the four QN-rich regions within the PD of Rnq1 showed that while any two were generally sufficient for the maintenance of the [*PIN^+^*] prion, deletions of individual QN-rich regions created transmission barriers, and the possibility of transmission across these barriers depended on the presence of at least one common QN-rich region between the donor and recipient constructs. This underscored both the importance of identical amino-acid stretches within the PD for prion transmission and the contribution of conformational differences created by other parts of the PD (Kadnar et al., 2010). Our current study contributes to understanding exclusively conformation-based transmission barriers because the sequence of the entire PD is identical for Rnq1_WT_ and Rnq1_T27P_.

We found no evidence that the initial transmission of the prion state from [*PIN^+^_WT_*] to Rnq1_T27P_ was blocked. Rnq1_T27P_ joins [*PIN^+^_WT_*] amyloid aggregates as efficiently as Rnq1_WT_ and does not “poison” them (Figs 6A, 6B, 8A, 8B, and S2). But when we scored individual [*PIN^+^_T27P_*] isolates from post-transmission cultures for prion maintenance, the striking observation is that all post-transmission [*PIN^+^_T27P_*] isolates were extremely unstable. Scoring lineages of [*PIN^+^_T27P_*] isolates for prion maintenance following a controlled number of mitotic divisions between rounds of analysis revealed that, while on average mitotic stability of [*PIN^+^_T27P_*] increased compared to the first round of analysis (Fig 5C), mitotic growth also led to the establishment of [*PIN^+^_T27P_*] variants with distinct mitotic stabilities. Along with highly stable [*PIN^+^_T27P_*]s (>99.9% maintenance after 19 cell divisions) we obtained lineages with a heritable stability of ∼20% (Fig 5D). Importantly, like [*PIN^+^_WT_*] (Derkatch et al., 2000), [*PIN^+^_T27P_*] did not provide a detectable growth advantage relative to [*pin^-^_T27P_*], so there was no selection for cells with more efficiently propagating [*PIN^+^_T27P_*]. Thus, mitotic stability is just a quantitative characteristic that can be used to distinguish different [*PIN^+^_T27P_*] variants established after crossing the transmission barrier.

Two models could explain the appearance of heritable [*PIN^+^_T27P_*] variants after multiple mitotic divisions in the presence of [*PIN^+^_WT_*] (Fig 9). Both postulate that the T27P mutation puts conformational constraints on the PD of Rnq1_T27P_, so it cannot form a stable [*PIN^+^*] that is conformationally similar to [*PIN^+^_WT_*] even though Rnq1_T27P_ can form stable prions *de novo*. The **first model** proposes that multiple [*PIN^+^_T27P_*] variants are formed in the same cell after transmission, and then daughter cells inherit them randomly, with a higher chance of inheriting variants that produce more seeds (propagons) and are thus more stable in mitotic divisions. This would lead to the rapid disappearance of the least stable variants in the population. However, following individual [*PIN^+^_T27P_*] pedigrees allowed us to detect such unstable variants. In support of this model, the co-existence of different prion variants in the same cell for several generations has been confirmed for [*PSI^+^*] (Norton et al., 2024). The **second model** predicts that the transmission barrier leads to formation of a particular [*PIN^+^_T27P_*] variant, but this variant is not conformationally stable and undergoes frequent conformational switches until more energetically favorable states are attained. These stable conformations may have different, even low, mitotic stabilities, but they are not changing frequently anymore. The process of such conformational adaptation may involve multiple steps and some [*PIN^+^_T27P_*] variants may end up in efficiently propagating but “undecided” states that have no preferred path towards conformational stabilization. Our detection of such “undecided” [*PIN^+^_T27P_*] lineages that produce both more and less mitotically stable variants even after 100 cell divisions (Fig 5D and data not shown) supports the conformational adaptation model, but we do not see the models as strictly mutually exclusive. Conformational adaptation in the absence of PD mutations has been proposed e.g. for *de novo* induced [*PIN^+^*] and [*PSI^+^*], for permanent [*PSI^+^*] variant switching in different chaperone environments, and for conversion of *in vitro* made PrP^Res^ into PrP^Sc^ (Derkatch et al., 2001; Sharma and Liebman, 2012; Baskakov, 2014; King, 2022).

**Fig 9.**
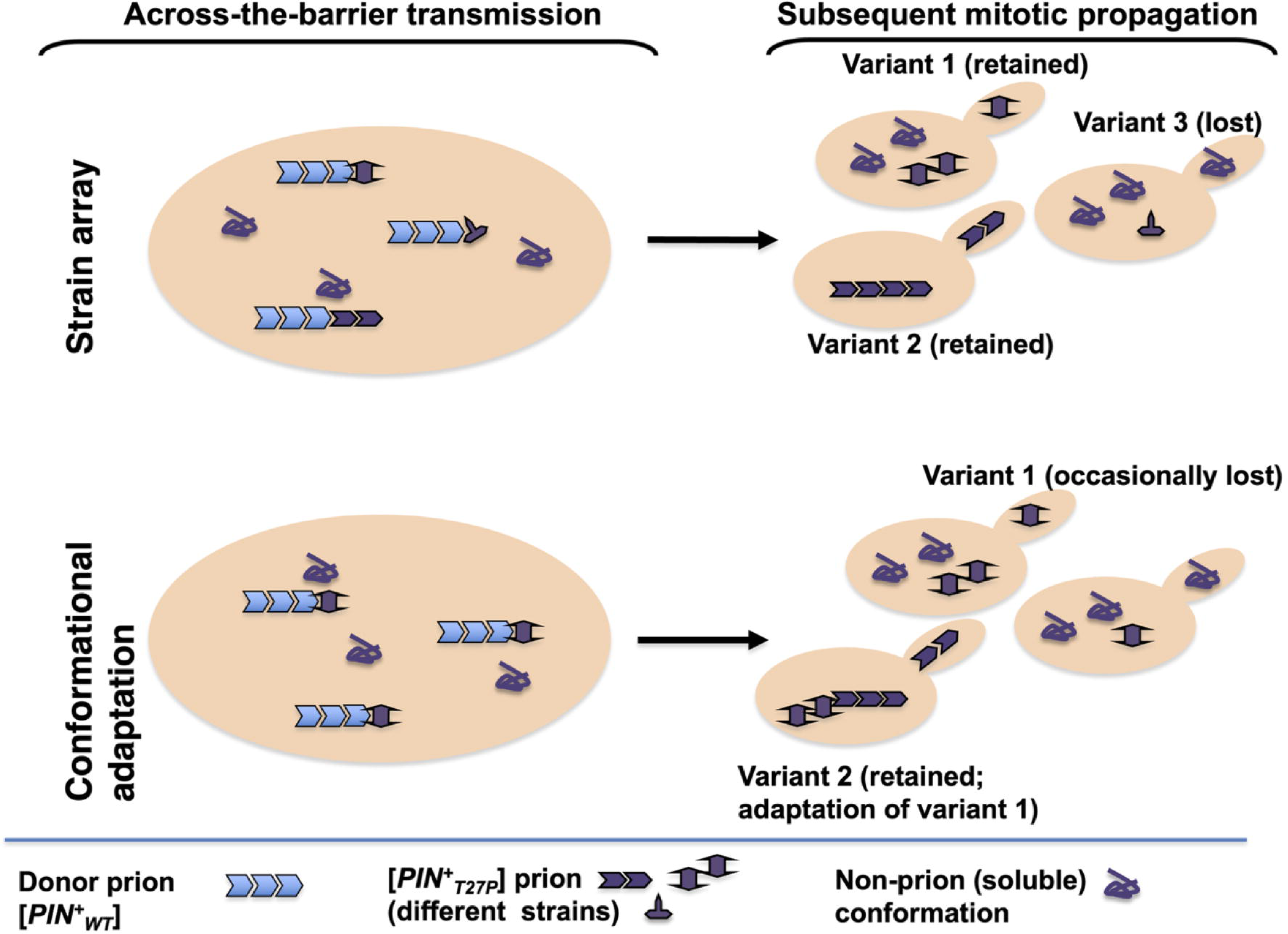
Models explaining the establishment of heritable [*PIN^+^_T27P_*] variants after across-the-barrier transmissions from [*PIN^+^_WT_*]. See text.

The T27P mutation specifically interacts with the B1C2 part of the PD since its deletion dramatically increased prion transmission to mutant constructs (Fig 7). Actually, data indicate that there is no transmission barrier between [*PIN^+^_τιB1C2-WT_*] and Rnq1_τιB1C2-T27P_. Even though the mitotic stability of [*PIN^+^_τιB1C2-T27P_*] was not very high, this level of stability was clearly heritable during subsequent mitotic growth, indicative of the establishment of a [*PIN^+^_τιB1C2-T27P_*] variant not undergoing further conformational adaptation (Fig 7C). The alternative explanation that a prion variant used as a [*PIN^+^_τιB1C2-WT_*] donor is more compatible with prion conformations that could be attained by the T27P mutants is unlikely since deletion of B1C2 also affects the transmission from [*PIN^+^_FL-WT_*] to the Rnq1_τιB1C2-T27P_ mutant (Fig 7D). While initially the barrier was strong, mitotic stability of the [*PIN^+^_τιB1C2-T27P_*] isolates increased much faster compared to [*PIN^+^_FL-T27P_*]s obtained from the same donor. Thus, despite carrying the same mutation, Rnq1_FL-T27P_ and Rnq1_τιB1C2-T27P_ form different prion variants. Analysis of transmissions between Rnq1 fragments carrying deletions of only QN1 (ΔB1) or QN2 (Δ2D) regions indicates that simultaneous deletion of both regions is required for the lifting of the transmission barrier created by the T27P mutation, but removing just B1 encompassing a 12 aa QN-rich region is sufficient to reduce the strength of the barrier.

Another indication of a difference in amyloid conformations attained by Rnq1_WT_ and Rnq1_T27P_ is that amyloid fibers spontaneously formed by these proteins *in vitro* look different in TEM: the bundles of Rnq1_WT_ fibrils are tight and twisted whereas Rnq1_T27P_ fibers are more relaxed and not twisted (Fig 4C).

However, the T27P mutation affects more than the array of prion conformations that are possible for the [*PIN^+^_T27P_*] prion. It changes the conformation of the protein even in the soluble state, indicated by a slight but reproducible increase of mobility of SDS-PAGE gels when purified from yeast or bacteria (Figs 2A and 4A). Indeed, T27P opens an alternative aggregation pathway for the Rnq1 protein. The Rnq1_WT,_ even when overexpressed, is not prone to aggregate when fused to C-terminal GFP family tags unless a heterologous QN-rich amyloid aggregate is present (Derkatch et al., 2001). This may be because a bulky tag blocks QN4, the last QN-rich determinant in the PD, which has recently been shown to initiate [*PIN^+^*] formation (Galliamov et al., 2024, 2025), so heterologous amyloid seeds are needed to overcome this constraint. But Rnq1_T27P_ attached to YFP readily forms liquid-like droplets at physiological levels of expression without the need for heterologous prions (Fig 8). This finding strengthens the evidence that the T27P mutation affects aggregation properties of Rnq1.

### Is the only function of the Rnq1 “NPD” to regulate the aggregation of Rnq1 PD?

Transmission barriers created by mutations **within the PDs** of yeast prions (see above) as well as the existence of different prion variants can be explained by the parallel-in-register model of protein molecules in amyloid fibrils (Wickner et al., 2008; 2011a). According to this model, prion variants are the result of differences in the length of amyloid cores formed by PDs and relative positioning of amyloid stretches due to turns within regions not involved in β-strand formation. For Rnq1, amyloid formation was detected in the regions roughly corresponding to QN-rich regions, but varies in different prion variants (Galliamov et al., 2024, 2025). So, mutations within or deletions of these regions are expected to create conformational constrains leading to different prion variants. But the T27P mutation is located **outside** the currently defined Rnq1 PD, so its effect on variants of the Rnq1-based prions is not well explained by this model.

How could the T27P mutation in the NPD create a transmission barrier? Because NPDs engage in interactions with cellular partners, the absence or inactivation of NPD can block these interactions and thereby increase *de novo* prion formation (Derkatch et al., 1996; Maddelein and Wickner, 1999). Increasing such interactions would, reciprocally, decrease prion formation (Derkatch et al., 1998). However, if the prion-forming protein is indeed chimeric and the NPD and PD domains fold independently, mutations that just change functionality of the NPD are not expected to impose **specific** conformational constrains on the PD and should not lead to transmission barriers. An alternative hypothesis is that in Rnq1 the N-terminal domain, presumed NPD, regulates the conformation and aggregation properties of the PD.

Indeed, this hypothesis helps explain multiple previous findings indicating that the NPD of Rnq1 is involved in essentially all prion-related phenotypes associated with [*PIN^+^*]. First, while constructs where Rnq1 PD is separated from Rnq1 NPD and attached to Sup35MC can form prions, these prions have distinct properties from [*PIN^+^*] formed by Rnq1_FL_ (Sondheimer and Lindquist, 2000; Bardill and True, 2009). Second, *in vitro* Rnq1_FL_ forms amyloid considerably slower than Rnq1 fragments lacking the NPD (Vitrenko et al., 2007b; Kadnar et al., 2010), which is consistent with inhibition of aggregation by the NPD even in the absence of its cellular interactors. Third, point mutations and deletions in Rnq1 NPD affect the propagation of [*PIN^+^*] (Vitrenko et al., 2007a, Shibata et al., 2009; Kurahashi et al., 2011a). Curiously, among 13 missense mutations uncovered by a screen for loss of [*PIN^+^*], 12 were in the NPD and two mutations, V23A and A29T, were located very close to our T27P mutation, within the same predicted helical stretch (Shibata et al., 2009; Kurahashi et al., 2011a). The authors interpreted their results as moderate destabilization of [*PIN^+^*] by the mutations and the possibility of transmission barrier was not tested. However, our results suggest that data in these earlier papers are also consistent with the creation of transmission barriers between [*PIN^+^_WT_*] and at least some of the mutants. The Shibata et al. study also supports the likelihood that our genetic screen was close to saturating and that mutations creating strong transmission barriers, like T27P, are extremely rare while most other changes have much milder effect (between 30 and 70% of cells carried [*PIN^+^_MUTANT_*] prions in their experiments corresponding to Round 1 analysis here). Fourth, a site for interaction with the Sis1 chaperone essential for [*PIN^+^*] propagation is located in the NPD (aa 91-97) and mutations in this site lead to [*PIN^+^*] destabilization (Douglas et al., 2008; Shibata et al., 2009). Overexpression of Rnq1_FL_, but not NPD or PD alone, is toxic in [*PIN^+^*] but not [*pin^-^*] cells due to oxidative stress, and this toxicity is rescued by overexpression of the Sis1 chaperone (Douglas et al., 2008; Bharthi et al., 2016). To explain this, it was hypothesized that Sis1 directs Rnq1 into non-toxic [*PIN^+^*] amyloid aggregates, preventing it from forming off-pathway SDS-soluble aggregates that cause oxidative stress (Douglas et al., 2008). Finally, the NPD of Rnq1 also affects interactions of [*PIN^+^*] with other prions. Overexpression of a fragment with a specific deletion in NPD, Rnq1_τι1-100_, leads to loss of [*PSI^+^*], [*URE3*] and huntingtin polyQ aggregates in [*PIN^+^*] but not in [*pin^-^*] cells (Kurahashi et al., 2008). Later, similar [*PIN^+^*]-dependent curing of [*PSI^+^*] was reported for multiple missense alleles in the Rnq1 NPD, including L26S, S28P and A29T located close to T27P (Kurahashi et al., 2009, 2011a, 2011b). One possible explanation based on our study is that these mutant proteins switch to variant(s) of [*PIN^+^*] that are incompatible with [*PSI^+^*], like those reported earlier (Bradley and Liebman, 2003; Mathur et al., 2009; Villali et al., 2020).

Curiously, several studies mention that changes in the NPD of Rnq1 led it to form non-amyloid aggregates when tagged with YFP, but these aggregates are different from Rnq1_T27P_ liquid-like droplets detected in our study. The SDS-soluble aggregates formed by Rnq1 with a mutated Sis1 binding site are highly toxic even in [*pin^-^*] cells (Douglas et al., 2008), while Rnq1_T27P_-YFP droplets do not inhibit growth. Also ThT-negative aggregates formed by Rnq1_τι1-100_-YFP (Kurahashi et al., 2008) actively recruit Rnq1_FL-WT_-CFP which does not join liquid-like droplets formed by T27P. Thus, the Rnq1 NPD can probably control formation of not one, but multiple types of non-amyloid aggregates.

While the NPD of Rnq1 is associated with multiple aggregation-related phenotypes, there is no evidence for its activity not related to Rnq1 aggregation. The *RNQ1* gene is not essential, and its deletion or overexpression does not lead to any significant growth defects in [*pin^-^*] cells (www.yeastgenome.org and our data).

The key rationale for [*PSI^+^*], [*URE3*], and [*PIN^+^*] being diseases rather than functional prions is based on their lower-than-expected frequency in wild yeast populations (Wickner et al., 2011b; Kelly and Wickner, 2013). But [*PIN^+^*] is detected in wild and commercial yeast isolates much more frequently than [*PSI^+^*] and [*URE3*], >10% *vs* <1% (Resende et al., 2003; Nakayashiki et al., 2005; Halfman et al.,2012; Westergard and True, 2014a; Kelly et al., 2014). So, the evidence for its detrimental effect is based on careful assessment of the frequency of outcross matings expected to spread prions and on polymorphism of the PD-encoding part of *RNQ1* locus expected to create transmission barriers (Kelly et al., 2012; Kelly and Wickner, 2013). These calculations assume stable inheritance of [*PIN^+^*]. However, our study reveals [*PIN^+^*] variants with heritably low mitotic stability. In addition to conformational switches induced by a transmission barrier, unstable [*PIN^+^*]s also form *de novo* and during stress or otherwise changing environmental conditions (Derkatch et al., 2000; Newnam et al., 2011; Westergard and True 2014b), and their instability may be heritable. Also, meiotic stability of [*PIN^+^*] is not 100% (Bradley et al., 2002), and at least some of >40 different [*PIN^+^*] variants (Huang et al., 2013; Westergard and True, 2014a) may be even more meiotically unstable, like [*URE3*] and some [*PSI^+^*]s (Liebman and All-Robyn 1984; Wickner, 1994; Bradley et al., 2002). Finally, redundancy of the Rnq1 PD makes it resistant to transmission barriers created by mutations: both our genetic screen here and screens mentioned above uncovered NPD rather than PD mutations affecting [*PIN^+^*]. Thus, [*PIN^+^*] may not have a strong enough detrimental effect to be considered a disease.

In thinking about possible functions for the Rnq1 protein it is instructive to recognize that PDs of both Ure2 and Sup35 do contribute to cellular functions of these proteins (Hoshino et al., 1999; Hosoda et al., 2003; Shewmaker et al., 2007; Edskes et al., 2018), but they are not required for their well-known major jobs. For Sup35, the PD is not required for an essential role in translation termination, but it is critical for at least two non-essential “moonlighting” jobs. One involves interactions with the Pab1 protein (Hosoda et al., 2003), a major component of stress granules known to be enriched in aggregation-prone proteins contributing to their assembly (Fomicheva and Ross, 2021). The other “moonlighting job for Sup35 uncovered in our work (Li et al., 2014) involves formation of a two-protein amyloid composed of Sup35 PD and the QN-rich domain of Pub1. The [*PUB1 / SUP35*] complex associates with microtubules and promotes integrity of the cytoskeleton. Thus, while [*PSI^+^*] may indeed be an egoistic byproduct of Sup35 aggregation properties, Sup35 function requiring the PD depends on the ability to form prion-like aggregates. We hypothesize that the same is true for Rnq1. Its cellular function may involve aggregation through the PD, and the NPD’s role is in regulating such aggregation. One such possible function is positive and negative interactions with other aggregating proteins in order to induce or eliminate heterologous protein aggregates. Whether this role is carried out by [*PIN^+^*] or through other single- or multiprotein aggregates encompassing Rnq1 has yet to be established.

## Materials and Methods

Most of the genetic and biochemical methods used for analysis of the [*PIN^+^*] prion in yeast, especially those used in our laboratory, are described in a review by Liebman et al. (2006).

### Strains and Plasmids

#### Bacterial Strains

Unless specifically mentioned, strain DH5α was used (F^−^ φ80*lac*Z*ΔM15 Δ*(*lacZYA*-*argF*)*U169 recA1 endA1 hsdR17*(r_K_^−^, m_K_^+^) *phoA supE44* λ^−^*thi*-1 *gyrA96 relA1*). Strain KC8 was used to select for the yeast *HIS3* and *LEU2* markers when transformed with DNA isolated from cells harboring more than one plasmid (*pyrF::Tn5, hsdR, leuB600, trpC9830, lacX74, strA, galUK, hisB436*). Strain BL21-AI was used for recombinant protein purification (F^−^*omp*T *hsd*S_B_ (r_B_^−^, m_B_^−^) *gal dcm ara*B*::T7RNAP-tetA*).

#### Yeast Strains

The original [*PIN^+^*][*psi^-^*] isolate of the yeast 74-D694 yeast strain (1Y1, aka L1749; aka “high multi-dot” [*PIN^+^*]; Derkatch et al., 1997; Bradley et al., 2002) and its derivatives were used in all experiments. The genotype of 74-D694 is *MAT***a** *ade1-14* (UGA) *leu2-3,112 his3-Δ200 trp1-289* (UAG) *ura3-52* (Chernoff et al., 1995). The 74-D694-based [*PIN^+^*][*psi^-^*] *rnq1-Δ* strain, YID108, and independently obtained isolates of this strain, YID146.2 and 146.3, were constructed as described in Kadnar et al. (2010). In these strains the entire *RNQ1* ORF and ∼100 bp *RNQ1* terminator sequence is seamlessly disrupted, and [*PIN^+^*] is maintained by the *CEN URA3* plasmid carrying the *RNQ1_WT_* ORF driven by the native *RNQ1* promoter and followed by the native *RNQ1* terminator sequence (RNQ1_WT;_ pID127). We have previously confirmed (Kadnar et al., 2010) that in these strains Rnq1 is expressed at a level similar to that of endogenous Rnq1 and [*PIN^+^*] is stably maintained (>99.5% stability). Unless specifically mentioned, the YID108, YID146.2, and YID146.3 strains also carry the *CEN HIS3 GAL-SUP35NM-YFP* plasmid (pID106; Kadnar et al., 2010) allowing the presence of [*PIN^+^*] to be scored through the *de novo* [*PSI^+^*] induction. When indicated, plasmids pID349 (ΔB1C2), pID341 (ΔB1), pID342 (Δ2D), and pID344 (Δ3E) carrying *RNQ1_WT_* fragments were substituted for the full-length *RNQ1_WT_* as [*PIN^+^*] maintainers in the YID146.3 strain (see Plasmids). We have previously demonstrated that these fragments are expressed at their expected sizes and can stably maintain the [*mini-PIN^+^*] prions and promote [*PSI^+^*] induction (Kadnar et al., 2010). Stable [*mini-PIN^+^*] isolates with these maintainers, YID412, YID405, YID407, and YID409, respectively, are from the Kadnar et al. (2010) study.

A [*pin^-^*][*psi^-^*] *rnq1-Δ* strain (YID193) used for analysis of the *de novo* [*PIN^+^*] induction by overexpression of Rnq1_T27P_ and analysis of the *de novo* aggregation of Rnq1_T27P_-YFP in the *rnq1-Δ* background was obtained by selectively eliminating the *URA3* [*PIN^+^*] maintainer from YID146.3 on FOA media while maintaining the *GAL-SUP3NM-YFP* [*PSI^+^*] inducing plasmid. The [*pin^-^*][*psi^-^*] isolate of 74-D694 (1G4; Derkatch et al., 1997) obtained by curing the [*PIN^+^*] prion on media containing 5mM GuHCl was used in experiments testing the *de novo* aggregation of Rnq1_T27P_-YFP and Rnq1_WT_-YFP in the presence of chromosomally encoded Rnq1_WT_.

### Plasmids

All yeast plasmids used in this study include the same modules: promoters are *Eco*RI – *Bam*HI fragments, ORFs are *Bam*HI – *Sac*II fragments, and terminators / fluorescent tags are *Sac*II – *Sac*I fragments inserted into the poly-linkers of the pRS series plasmids (Sikorski and Hieter, 1989). Site-directed mutagenesis was done using the QuickChange kit (Stratagene) according to manufacturer’s instructions and primers #360 and #361 (see Table S2).

The pRS416-based *CEN URA3* (pID127) and the pRS415-based *LEU2 CEN* (pID129) plasmids carrying the *RNQ1_WT_* ORF driven by the native *RNQ1* promoter and followed by the native *RNQ1* terminator sequence (*RNQ1_WT_*) are described in Kadnar et al. (2010). To make these plasmids, the *RNQ1* ORF was PCR amplified from 74-D694 chromosomal DNA. An earlier study (Kadnar et al., 2010) confirmed that from these constructs Rnq1_WT_ is expressed at a level similar to that of the chromosomally encoded Rnq1 which allows for stable maintenance of [*PIN^+^*] and does not lead to the induction of the *de novo* appearance of [*PIN^+^*] in *[pin^-^*] cells.

The *RNQ1_T27P_* carrying plasmid and other plasmids in the library used for the genetic screen are pRS415-based, i.e. *LEU2 CEN*, and are identical to pID129 except for the mutagenized *RNQ1* ORFs. The *RNQ1_T27P_* carrying pID335 plasmid was obtained by site-directed mutagenesis of the pID219 plasmid.

Plasmids carrying the T27P mutation within *RNQ1* fragments lacking the sequences encoding the indicated QN regions and adjacent hydrophobic helices within Rnq1 PD were constructed by site-directed mutagenesis introducing T27P into the constructs previously described in Kadnar et al. (2010): pID331 encoding Rnq1_τιB1C2-T27P_ is based on the Rnq1_τιB1C2-WT_ (pID370; ΔB1C2) plasmid lacking QN1, QN2 and helices B and C; pID332 encoding Rnq1_τιB1-T27P_ – on the Rnq1_τιB1-WT_ (pID369; ΔB1) plasmid lacking QN1 and helix B); pID333 encoding Rnq1_τι2D-T27P_ – on the Rnq1_τι2D-WT_ (pID358; Δ2D) plasmid lacking QN2 and helix D; and pID334 encoding Rnq1_τι3E-T27P_ – on the Rnq1_τι3E-WT_ (pID387; Δ3E) plasmid lacking QN3 and helix E (Fig 1A). The pID370, pID369, pID358, and pID387 plasmids were used as controls for experiments with the corresponding abovementioned *LEU2* constructs carrying *RNQ1*_T27P_. Except for deletions in the *RNQ1* PD these plasmids are identical to pID335 (RNQ_FL-T27P_) and pID129 (RNQ_FL-WT_), respectively: pRS415-based, *LEU2 CEN,* original *RNQ1* promoter and terminator. Plasmids pID349 (Rnq1_τιB1C2-WT_), pID341 (Rnq1_τιB1-WT_), pID342 (Rnq1_τι2D -WT_), and pID344 (Rnq1_τι3E-WT_) are identical to pID370, pID369, pID358, and pID387, respectively, except that, like pID127, they carry the *URA3* marker. These plasmids were used as [*PIN^+^*] maintainers for experiments testing transmission barriers to Rnq1_FL-T27P_ or Rnq1 fragments carrying Rnq1_T27P_.

To overexpress Rnq1_WT_ and Rnq1_T27P_, respectively, the pID336 and pID337 plasmids were constructed. Both plasmids are pRS416-based, *URA3*, *CEN* and the full-length ORFs are controlled by the *CUP1* promoter and followed by the original *RNQ1* terminator.

In the pRS416-based *CEN LEU2 CUP-RNQ_WT_-CFP* plasmid (pID121; Kadnar et al., 2010), a fusion of complete *RNQ1_WT_* ORF to *CFP*, is driven by the inducible *CUP1* promoter. The pRS415-based *URA3*-marked pID403 and pID404 plasmids are similar except they carry, respectively, *CUP-RNQ_WT_-YFP* and *CUP-RNQ_T27P_-YFP*.

The pRS413-based *CEN HIS3 GAL-SUP35NM-YFP* plasmid (pID106, Kadnar et al., 2010) carries *SUP35NM-YFP* encoding the PD of the [*PSI^+^*]-forming Sup35 protein fused to the YFP reporter and placed under control of the inducible *GAL1* promoter.

Bacterial expression plasmids for Rnq1_WT_ and Rnq1_T27P_ are pJC45-based and express N-terminally 10xHIS tagged proteins. The ΔB1C2D3_WT_ plasmid (pID257; aka QN4 only) and full-length RNQ1_WT_ (pID249/250) constructs are described in Kadnar et al. (2010) and Vitrenko et al. (2007b), respectively. The ΔB1C2D3_WT_ construct expresses the Rnq1NPD through the QG repeat region (aa1-172) fused to the helical region E and QN4 in the PD (aa 320-405); the fusion is seamless, *i.e* no additional amino acids are introduced in between. The ΔB1C2D3_T27P_ and RNQ1_T27P_ differ only by the T27P mutation that was introduced by site-directed mutagenesis into ΔB1C2D3_WT_ and RNQ1_WT_, respectively.

### Yeast Media and Cultivation

Standard yeast media and cultivation procedures were used (Sherman et al., 1987; Rose et al., 1990; Amberg et al., 2005). Unless specifically mentioned yeast transformants were grown at 30^0^C on solid synthetic glucose media selective for plasmid maintenance (e.g. SD-UraLeuHis). In addition to synthetic glucose SD-Ade media, synthetic ethanol SEt-Ade media supplemented with 2% ethanol as the only carbon source was used to score for [*PSI^+^*] (see below). Untransformed strains and cultures for protein isolation were grown on complete glucose media, YPD. The *GAL* promoter was induced on solid synthetic media selective for plasmid maintenance and supplemented with 2% galactose as a single carbon source (e.g. SGal-LeuHis). The *CUP* promoter was induced by supplementing SD media selective for plasmid maintenance with the indicated concentration of CuSO_4_ (5, 20 or 50μM; see Results and Figure Legends for details of specific experiments). To selectively eliminate plasmids with the *URA3* marker, yeast were passaged twice on solid SD supplemented with 5-fluoroorotic acid (5-FOA; Boeke et al., 1984) selective for other plasmids. Solid YPD supplemented with 5mM guanidine hydrochloride (GuHCl; Tuite et al., 1981) was used to cure the [*PSI^+^*] and [*PIN^+^*] prions. YPGlycerol media supplemented with 2% glycerol as a single carbon source was used to detect Pet^-^ colonies that may be confused with [*PSI^+^*] by color.

### *RNQ1* Mutagenesis and Genetic Screen

To mutagenize *RNQ1*, the complete 1.2 kb *RNQ1* ORF was PCR amplified from pID129 using *Taq* polymerase (New England Biolabs), primers # 7 and # 8 (see Table S2), and manufacturer’s recommended reaction conditions, which was expected to yield ∼1 mutation per 1,000 bps after 32 cycles. To eliminate mutagenic bias from any one PCR reaction, seven independent PCR reactions were pooled. PCR products were digested with *Bam*HI and *Sac*II and re-introduced into the *Bam*HI- and *Sac*II- digested pID129 vector between the *RNQ1* promoter and *RNQ1* terminator sequences with a stop codon right after the *Sac*II site. Ligation reactions were transformed into *E. coli*. Approximately 30,600 transformant colonies were washed off with ∼1 ml LB from three transformant selection plates (banks pID142 – pID144) and ∼200μl of suspensions were added to 50ml LB+Amp (30 mg/l) and grown for five hours at 37^0^C prior to plasmid DNA isolation. Plasmid DNA was introduced into the YID108 yeast cells. Sequencing the *RNQ1* ORF from two randomly selected *E. coli* transformants with primers #119 and #8 confirmed the presence of 1 mutation per the Rnq1 PD. At this estimated rate of mutagenesis, screening ∼12,000 plasmids was expected to result in >95% chance of scoring all three possible nucleotide changes at each base.

The genetic screen is described in Results and is illustrated in Fig 1B. Approximately 40 patches of yeast transformants were analyzed per plate. Each plate had negative (EV) and positive (pID129) control transformants. Passages on all indicated media were by replica plating. Two passages on SD-LeuHis not selective for the <u>*URA3 RNQ1_WT_* plasmid preceded replica plating to FOA to accumulate cells that lost this plasmid.</u> Two passages on 5-FOA were sufficient for the selective elimination of all cells carrying the *URA3*-marked *RNQ1_WT_* plasmid (confirmed by replica plating to SD media lacking uracil). Induction of [*PSI^+^*] was scored, after one and two passages on SGal-LeuHis, on SD-Ade at 30^0^C and 20^0^C and on SEt-Ade at 30^0^C after up to 25 days of incubation, which is sufficient for the detection of even the weakest [*PSI^+^*] variants.

No [*PSI^+^*] induction was detected in ∼500 transformants with mutated plasmids. Immediately excluded were the candidates that: (i) were Pet^-^ (detected as lack of growth on YPGlycerol to which all patches were also replica plated); (ii) carried *RNQ1_MUTANT_* plasmids lacking the full-length 1.2 kb *RNQ1* ORF insert (detected by PCR using primers #7 and #8 and total yeast DNA from after-shuffle cultures). Also, to exclude plasmids that carried frameshift mutations (usually -1) within the forward primer used to amplify the *RNQ1* ORF or stop codons in the very beginning of the ORF, a blue-white ligation-based assay was developed. The first ∼100 bps of the *RNQ1* ORF were PCR amplified using forward primers #127 or #124 and reverse primer #129 and cloned into the pUC18 plasmid as *Bcl*I – *Hin*dIII, *Bam*HI – *Hin*dIII, or *Bgl*II – *Hin*dIII fragments to create the in-frame, -1, and +1 fusions with the *lacZ* gene. The ligation mixes were transformed into the DH5α *E. coli* strain to test for the complementation of the φ80*lac*ZΔM15 allele. White or light blue color of most bacterial transformants on the LB+Amp media supplemented with 40μg/ml XGal was indicative of the disruption of LacZ synthesis in this particular frame. The test was validated by sequencing ∼30 plasmids with presumptive early frameshift and nonsense mutations and >100 plasmids with no mutations in the beginning of the *RNQ1* ORF predicted by this assay.

For the remaining candidates, the *RNQ1_MUTANT_* plasmids were isolated for further analysis. To selectively isolate only the *LEU2-*marked *RNQ1_MUTANT_* plasmids and not the *HIS3*-marked *GAL-SUP35NM-YFP* plasmids, the *E. coli* KC8 strain (see *Strains* above) was transformed with total yeast DNA extracts, and Leu^+^ transformants were selected on M9-based media lacking leucine. Digestion of plasmids with restriction endonucleases was used to exclude *RNQ1_MUTANT_* plasmids with re-arrangements outside of the *RNQ1* ORF. Then, for each candidate, DNA from at least two independent *E. coli* transformants was re-transformed into the original yeast strain and re-tested according to the same scheme as in the original screen (see Fig 1B); in total, at least 12 yeast transformants were tested for each candidate. The re-transformation test provided proof that a mutation in the *RNQ1_MUTANT_* plasmid was the cause of the Pin^-^ phenotype in the after-shuffle cultures. Alternatively, the Pin^-^ phenotype could be either due to spontaneous loss of the [*PIN^+^*] prion (0.2 – 0.5% in control experiments where the *RNQ1_WT_* plasmid was shuffled-in) or due to chromosomal mutations e.g. either eliminating [*PIN^+^*] or inhibiting [*PSI^+^*] induction. Primers #7, #8, #17, #18, #19, and #30 were used to sequence the *RNQ1* ORF and adjacent promoter and terminator regions (see Table S2).

### Scoring for the presence of the [*PIN^+^*] prion and Pin phenotype

Induction of the *de novo* appearance of the [*PSI^+^*] prion upon overexpression of the PD of the Sup35 protein, Sup35NM, was used to score for the presence of [*PIN^+^*] (see Liebman et al., 2006). The assay is based on our finding that [*PIN^+^*] is required for the efficient induction of [*PSI^+^*] (Derkatch et al., 1997). In [*PIN^+^*] cells expressing Sup35NM-YFP at an ∼5-fold increased level compared to the expression of endogenous Sup35 for ∼10 cell generations, [*PSI^+^*] forms in ∼10% cells, whereas appearance of [*PSI^+^*] in [*pin^-^*] cultures is extremely rare, <10^-4^. Transformants carrying the *CEN HIS3 GAL-SUP35NM-YFP* [*PSI^+^*] inducing construct and other indicated plasmids were grown on SD media selective for all the plasmids and then replica plated onto the inducing SGal media, also selective for all the plasmids. Growth on SGal was for 2 days (5-7 generations), a second passage on SGal was done to increase the number of generations and thus the chance of [*PSI^+^*] induction. As a negative control, the same cultures were passaged on non-inducing SD media and then analyzed in parallel with induced cultures. Three methods were used to score for the appearance of [*PSI^+^*].

**(1)** For qualitative/semi-quantitative analysis, yeast were replica plated from SGal (or control SD) media onto SD media lacking adenine and not selective for plasmids. An Ade^+^ phenotype after SGal and not after SD is indicative of [*PSI^+^*] induction. Growth of [*PSI^+^*] cells on adenineless media is due to suppression of a premature UGA stop codon in the *ADE1* gene present in our strains (*ade1-14*). The nonsense suppression is due to the sequestration of the chromosomally encoded full-length Sup35 protein, a translation termination factor, into the [*PSI^+^*] prion aggregates initiated by Sup35NM-YFP overexpression. For all such experiments, both SD-Ade and SEt-Ade media were used. Cultures on SD-Ade were incubated at 30^0^C and 20^0^C; cultures on SEt-Ade were incubated at 30^0^C. Growth was recorded multiple times between 3 and 25 days of incubation. This approach allows detection of [*PSI^+^*] variants of different strengths (some very weak [*PSI^+^*] variants grow only on SEt-Ade or at lower temperatures, and only after a very long incubation).

**(2)** For quantitative analysis, yeast were colony purified on YPD from galactose (or control glucose) media. On YPD, [*PSI^+^*] cells form white or pink colonies whereas [*psi*^-^] cells are red due to the accumulation of a red pigment from the adenine precursor accumulating in the *ade1-14* cells. The appearance of white and pink Ade^+^ colonies (among red Ade^-^ colonies) following Sup35NM-YFP overexpression is indicative of [*PSI^+^*] induction.

However, nonsense suppression and white color in (1) and (2) can also be caused by genomic mutations in *SUP35* and other genes. Such genomic mutations are rare compared to the induced [*PSI^+^*] *de novo* formation in [*PIN^+^*] cultures. Yet, and especially in cultures where only a few Ade^+^ or white / pink colonies were detected, such colonies were confirmed to be [*PSI^+^*]. For the GuHCl test, white / pink colonies were patched onto YPD, then passaged three times on YPD+5mM GuHCl and then returned to YPD. Because GuHCl cures [*PSI^+^*] (Tuite et al., 1981), [*PSI^+^*] patches turn red on GuHCl media and remain red when returned to YPD. Chromosomal mutants may turn red on GuHCl (Bradley et al., 2003) but return to their white or pink color once back on YPD. Alternatively, for colonies that retained the *HIS3 GAL-SUP35NM-YFP* plasmid, the presence of [*PSI^+^*] was confirmed by fluorescence microscopy: after an overnight growth on SGal-His media, dot-shaped foci are detected in most cells in [*PSI^+^*] cultures.

**(3)** To directly visualize newly forming [*PSI^+^*]s, yeast cells from SGal media were analyzed by fluorescent microscopy utilizing the YFP tag attached to Sup35NM in the [*PSI^+^*] inducing construct. Newly forming [*PSI^+^*]s can be seen as ring-, line- or dot-shaped foci; cells with already established [*PSI^+^*] have dot foci; in [*psi^-^*] cells fluorescence is evenly distributed (Patino et al., 1996; Zhou et al., 2001).

### Assessing the [*PSI^+^*] prion variants

To assess the arrays of the [*PSI^+^*] prion variants induced in cells expressing Rnq1_T27P_ *vs* Rnq1_WT_, the Ade^+^ cells from patches replica plated from SGal to SD-Ade [*PSI^+^*] selective media and grown for 25 days at 20^0^C were thoroughly scraped and colony purified on YPD and individual colonies were randomly picked up from YPD for further analysis before color had developed. At least 12 transformants were analyzed for either Rnq1_T27P_ or Rnq1_WT_. The same approach was used to obtain [*PSI^+^*]s appearing in [*pin^-^*] cultures carrying EV. Alternatively, yeast were colony purified on YPD from the second pass on SGal-LeuHis media, where [*PSI^+^*]s were induced. Three assays were used to assess [*PSI^+^*] strength. **(1)** Color on YPD (the lighter the color, the stronger the [*PSI^+^*] variant). **(2)** Growth on adenineless media: SD-Ade at 30^0^C and at 20^0^C, and SEt-Ade at 30^0^C (the more robust the Ade^+^ phenotype, the stronger the [*PSI^+^*]). **(3)** Efficiency of curing by GuHCl: after three passages on YPD+GuHCl yeast were replica plated onto the same set of adenineless media as in (**2**). While weak [*PSI^+^*]s were usually cured completely, occasional Ade^+^ colonies were detected for stronger [*PSI^+^*]s.

### Analysis of the *de novo* induction of [*PIN^+^T27P*]

Both variants of this experiment are succinctly explained in Results related to Figs 3A and B. See Supplementary Materials and Methods for plasmid numbers and details on controls.

### Plasmid stability analysis

To compare stability of the Rnq1 expressing plasmids in yeast transformants carrying the *CEN URA3 RNQ1_WT_* and *CEN LEU2 RNQ1_T27P_* plasmids *vs* transformants carrying two *RNQ1_WT_* plasmids (*CEN URA3* and *CEN LEU2*) or a *RNQ1_WT_* (*CEN URA3*) plasmid and the EV (*CEN LEU2*), fresh transformants were patched onto SD-UraLeu media selective for both plasmids, then passaged twice on media selective for only one of the plasmids (SD-Ura or SD-Leu), then colony purified on the same media and replica plated to both that media and to the reciprocal media. Plasmid stability was determined as the fraction of colonies grown on the reciprocal media where only colonies formed by cells retaining the plasmid despite of the lack of selection were expected to grow.

### Western Blot Analysis of Rnq1T27P expression in yeast

Yeast cell lysates were prepared essentially as described in Liebman et al. (2006) and Kadnar et al. (2010). See Supplementary Materials and Methods for a detailed protocol. Either 30 or 60μg of total protein (concentration determined with BCA^TM^ Protein Assay Kit, Pierce) were separated on SDS-PAGE gels. Rnq1 was detected by Western blot with polyclonal antibodies raised in rabbits against the first 185 aa of Rnq1 (type 1A, a kind gift from E. Craig, University of Wisconsin-Madison). Sup35 was detected with a monoclonal antibody against the Sup35C raised in mouse (BE4). HRP-conjugated anti-rabbit and anti-mouse secondary antibodies and the Super Signal Western PICO ECL kit (Pierce) were used for detection.

### Recombinant protein purification from *E. coli*

Recombinant protein purification was essentially as described in Kadnar et al. (2010). See Supplementary Materials and Methods for a detailed protocol. Protein concentration was determined by the BCA^TM^ Protein Assay Kit (Pierce). Purity of recombinant proteins was estimated by SDS-PAGE as >95% (see Fig 4A).

### Analysis of *in vitro* fiber formation

Analysis of *in vitro* fiber formation by Rnq1_WT_ and Rnq1_T27P_ was performed essentially as in Kadnar et al. (2010). Specifically, 200 μl reactions were assembled in the fiber formation buffer (1M urea, final concentration; 100mM NaH_2_PO_4_, pH 7.4; 300mM NaCl) in the presence of 5 μM thioflavin T (ThT). Fiber formation was monitored by ThT fluorescence (LeVine, 1999) in a Molecular Devices SpectraMax M-5 plate reader at RT (ƛ_ex_=450 nm; ƛ_em_=483 nm). Readings were taken every 15 min, samples were shaken for 5 sec prior to each reading.

### Transmission Electron Microscopy (TEM)

TEM was performed at the Image Core Facility at the NYU School of Medicine. Following incubation to allow for amyloid fiber formation, protein / fiber suspensions were diluted 2.5-fold in water. Then 4 μl of diluted suspensions were applied to carbon-coated 400 mesh Cu/Rh grids (Ted Pella Inc.). Grids were washed 3 times with water to get rid of urea present in the fiber formation buffer. Then grids were negatively stained at 25^0^C with 1% uranyl acetate (applied twice briefly and then for 5 min). Images were captured using Philips CM12 TEM supplied with a Gatan 1k61k digital camera and processed using the Gatan Digital Micrograph software.

### Fluorescence Microscopy, ThT staining, and 1,6 Hexanediol treatment

Cells were observed using a Nikon Eclipse E600 microscope with a 60x / 1.4 NA oil immersion objective. YFP and CFP filters were used to observe YFP- and CFP-tagged constructs, respectively. Fluorescent and brightfield images of the same field were captured with a Zeiss AxioCam MRm digital camera and processed with AxioVision AxioVs40 V 4.8.2.0 software. In all experiments, representative groups of cells were scored, including cells of different sizes, with and without buds and with different levels of construct expression. For Rnq1 aggregation analysis, cells from at least three transformants were counted, at least 250 cells from each. For analysis of Sup35NM-YFP aggregation shown in Table 1 and Fig 2C, at least 11 vision fields were scored for each construct, ∼500 cells total per construct, while qualitative assessment of aggregation was done on a much larger scale. ImageJ software was used for cell counting. Photoshop was used for merging and adjusting images (only brightness and contrast were adjusted, no other processing). For all images shown, differences in cell size, budding and fluorescence intensity are within population variability range.

Staining with thioflavin T (ThT) was done essentially as described in Douglas et al. (2008). See Supplementary Materials and Methods for a detailed protocol. The CFP filter was used for ThT fluorescence analysis.

To dissolve liquid-like droplets with 1,6 Hexanediol (Hex), yeast transformants were grown overnight in liquid SD media selective for all plasmids. Then OD_595_ was adjusted to 1.0, CuSO_4_ was added to 5μM, and cultures were grown for two more hrs. Cells were harvested, resuspended in 100 – 500μl of the same SD media with 10% Hex (Fisher) and 0.2μg/μl digitonin (Sigma; 200 mg/ml stock in water) and incubated on a shaker for 10 min at 30^0^C. Then cells were centrifuged, resuspended in the same SD, and imaged using the filter matching the Rnq1 fluorescent tag.

### Statistics

Standard methods of statistical analysis were used. Averages and Standard Deviations (STDEV.S) were calculated in Excell (16.105.2 and earlier versions). Unless specifically mentioned, SD is included. Standard Error of the Mean (SEM) was calculated by dividing STDEV.S by square root of the number of datapoints. Excell and GraphPad Prism (10.6.1(892))were used for graphs. Statistics Kingdom (statisticskingdom.com) was used for T-tests (two sample Welch’s T-test; unknown / unequal SD). Unless specifically mentioned, the cutoff for the significance of variation was p<0.01.

## Abbreviations

NPD: non-prion domain
PD: prion domain
WT: wild type
GuHCl: guanidine hydrochloride
5-FOA: 5-fluoroorotic acid
Aa: amino acid
YFP: Yellow Fluorescent Protein
CFP: Cyan Fluorescent Protein
EV: empty vector
TEM: transmission electron microscopy
ThT: thioflavin T
Hex: 1,6 hexanediol
RT: room temperature
HFIP: 1,1,1,3,3,3 hexafluoroisopropanol

## Acknowledgements

We thank Elizabeth Craig for providing the anti-Rnq1 antibody. We are grateful to Gulnara Articov for help with *in-vitro* experiments and to Alice Liang and Eric Roth at the NYU School of Medicine Image Core Facility for performing TEM. We also thank Sei-Kyoung Park for helpful discussions of the data.

## Data availability

The authors confirm that all data underlying the findings are fully available without restriction. All relevant data are within the paper and its Supporting Information files.

## Author Contributions

Conceived and designed the experiments: ILD. Performed the experiments: SP, DMM, MLK, MA, APF, ILD. Analyzed the data: SP, MLK, SWL, ILD. Contributed reagents/materials/analysis tools: ILD, SWL. Wrote the manuscript: ILD, SWL.

## Supporting information

**Fig S1. The T27P mutation in Rnq1 does not lead to inhibition of growth in the presence of the highly expressed Sup35NM-YFP construct.** *RNQ1_WT_* was substituted with *RNQ1_T27P_* or control plasmids (indicated) in a [*PIN^+^*] [*psi^-^*] *rnq1-Δ* strain as described in Fig. 1B. Shown are the images of patches of five transformants for each plasmid substitution after they were replica plated from the *RNQ1_WT_* (*URA3*) counter-selecting FOA media to Sup35NM-YFP inducing SGal-LeuHis media or non-inducing SD-LeuHis. Before imaging, cells were incubated at 30^0^C for ∼48hrs. These are images of plates used for replica plating to adenineless media shown in Figure 2B.

**Fig S2. Rnq1_T27P_ joins [*PIN^+^_WT_*] prion aggregates as efficiently as Rnq1_WT_.** The [*PIN^+^*] 74-D694 strain was co-transformed, respectively, with the *CEN LEU2 CUP1-RNQ1_WT_-CFP* and *CEN URA3 CUP-RNQ1_T27P_-YFP* plasmids (**A**), or with *CEN LEU2 CUP1-RNQ1_WT_-CFP* and *CEN URA3 CUP-RNQ1_WT_-YFP* (**B**). Cultures of co-transformants were grown overnight (∼to mid-log) in no-copper SD-LeuHis media, diluted to OD_595_=1.0 into SD-LeuHis+5µM CuSO_4_ and grown for 2 more hours prior to observing the cells. In both images (A) and (B) CFP- and YFP-marked constructs co-localized in bright fluorescent foci in most cells (squares 1 and 2 for both (A) and (B); enlarged images are shown in panels below). However, there were cells with bright foci, for which only YFP or only CFP fluorescence is easily detectable on merged images. To test if there are cells in images from experiment (A) where only Rnq1_WT_ but not Rnq1_T27P_ joins [*PIN^+^_WT_*] aggregates, at total of ∼150 cells from three different transformants with only CFP detectable on merge images were analyzed more carefully. For all of them, either no evenly distributed YFP fluorescence was detected, indicative of the loss or rearrangement of the *CUP-RNQ1_T27P_-YFP* plasmid (see e.g. square 4 on Fig. S2A), or weak but definitely co-localizing YFP foci were detected on YFP images, indicative of reduced amounts of Rnq1_T27P_-YFP in the cell (e.g. square 5 on Fig. S2A). We also asked if there were Rnq1_T27P_-YFP aggregates not incorporating Rnq1_WT_. However, after analyzing ∼150 cells with only YFP detectable on the merge, we concluded that the lack of the CFP signal within these foci was also either due to the *CUP-RNQ1_WT_-CFP* plasmid loss (e.g. square 3 on Fig. S2A) or reduced abundance of Rnq1_WT_-CFP (e.g. square 6 on Fig. S6A). Analogous analysis in experiment (B) led to the same conclusion: squares 3 and 4 represent cells with reduced fluorescence for one of the constructs, and squares 5 and 6 – cells where one of the constructs is lost or rearranged.

**Fig S3. Growth on media supplemented with 5μM CuSO_4_ results in the level of Rnq1_T27P_-YFP expression similar to that of chromosomally encoded Rnq1_WT_.** A transformant of a [*pin^-^*] strain with the *CEN URA3 CUP-RNQ1_T27P_-YFP* plasmid was grown overnight (∼to mid-log) in the no-copper plasmid-selective SD-Ura media, then diluted to OD_595_=1.0 into SD-Ura+5μM CuSO_4_ and grown for 2 more hours prior to protein extraction (see Materials and Methods and Supplementary Materials and Methods). Approximately 30μg of total protein were separated on an SDS-PAGE gel. Rnq1 was detected by Western blot with polyclonal antibodies raised in rabbits against the first 185 aa of Rnq1 (type 1A Ab).

**Table S1. Similar arrays of [*PSI^+^*] variants are induced in cultures expressing Rnq1_T27P_, Rnq1_WT_, or not expressing Rnq1 after shuffling out the *CEN URA3 RNQ1_WT_* [*PIN^+^_WT_*] maintainer plasmid.** See Results and Materials and Methods for the description of the experiment. [*PSI^+^*]s isolated from at least eight independent transformants carrying the indicated plasmids were analyzed by the color test on YPD, growth on adenineless media and curability by GuHCl. Results of all three assays were consistent for ∼99% [*PSI^+^*] variants.

**Table S2. PCR primers used in this study.** Sites for restriction endonucleases used for cloning are underlined; the ATG start codon is in bold.

**File S1. Supplementary Materials and Methods.**

