## Supplementary figures and images for "Interactions between non-prion and prion domains of Rnq1 direct formation of amyloid vs liquid-like aggregates and create transmission barriers"

### Supplementary Figure 1

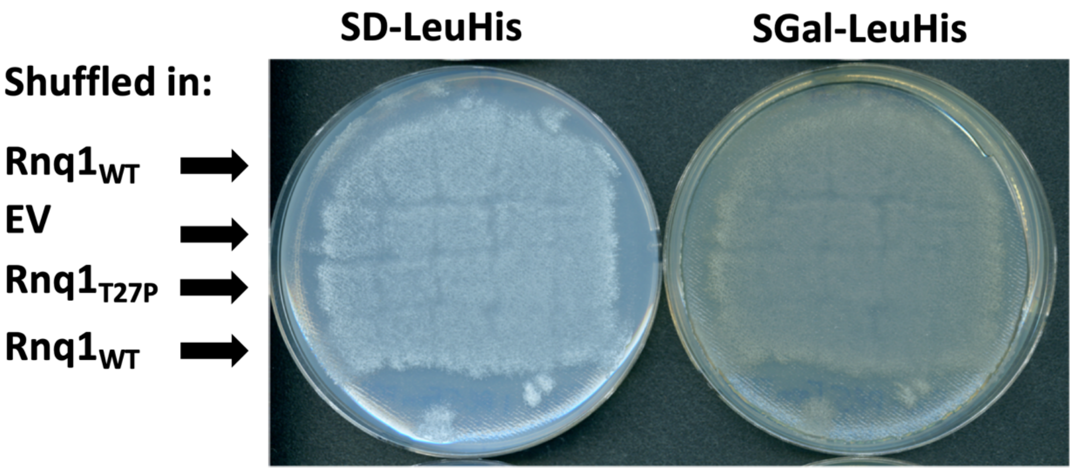

### Supplementary Figure 2

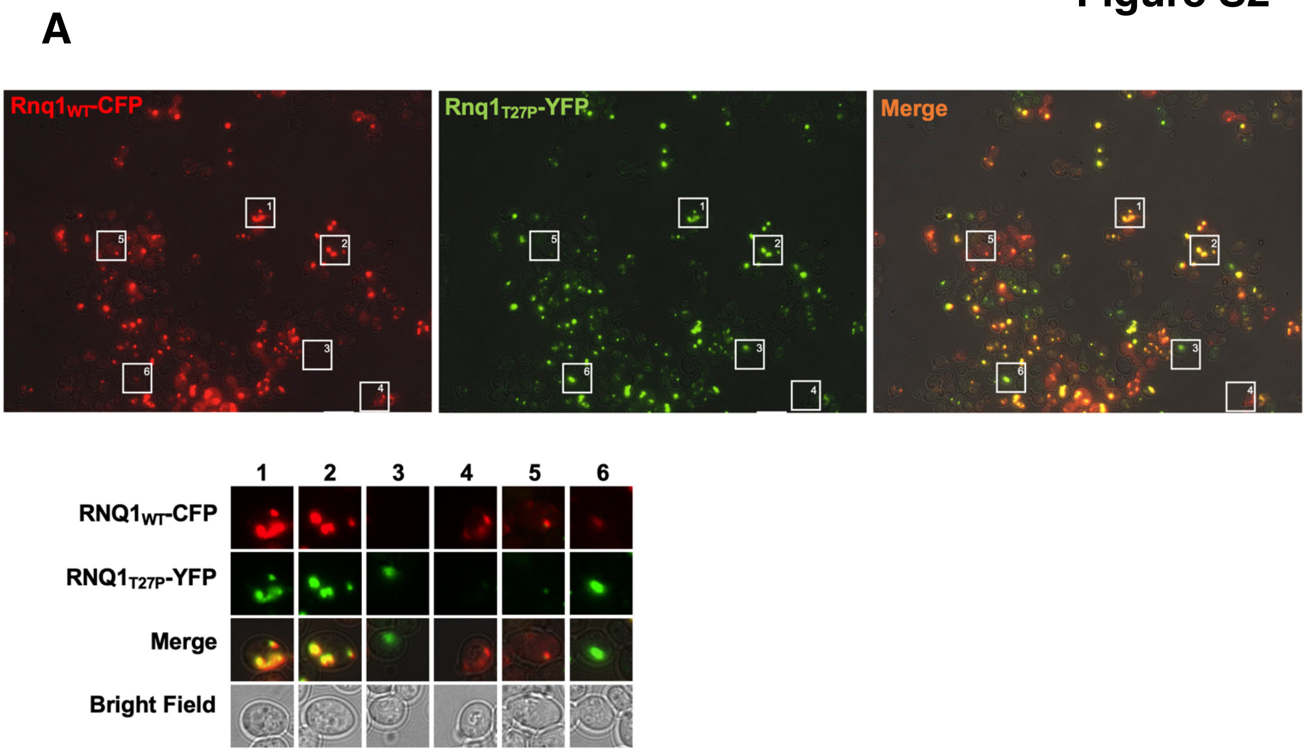

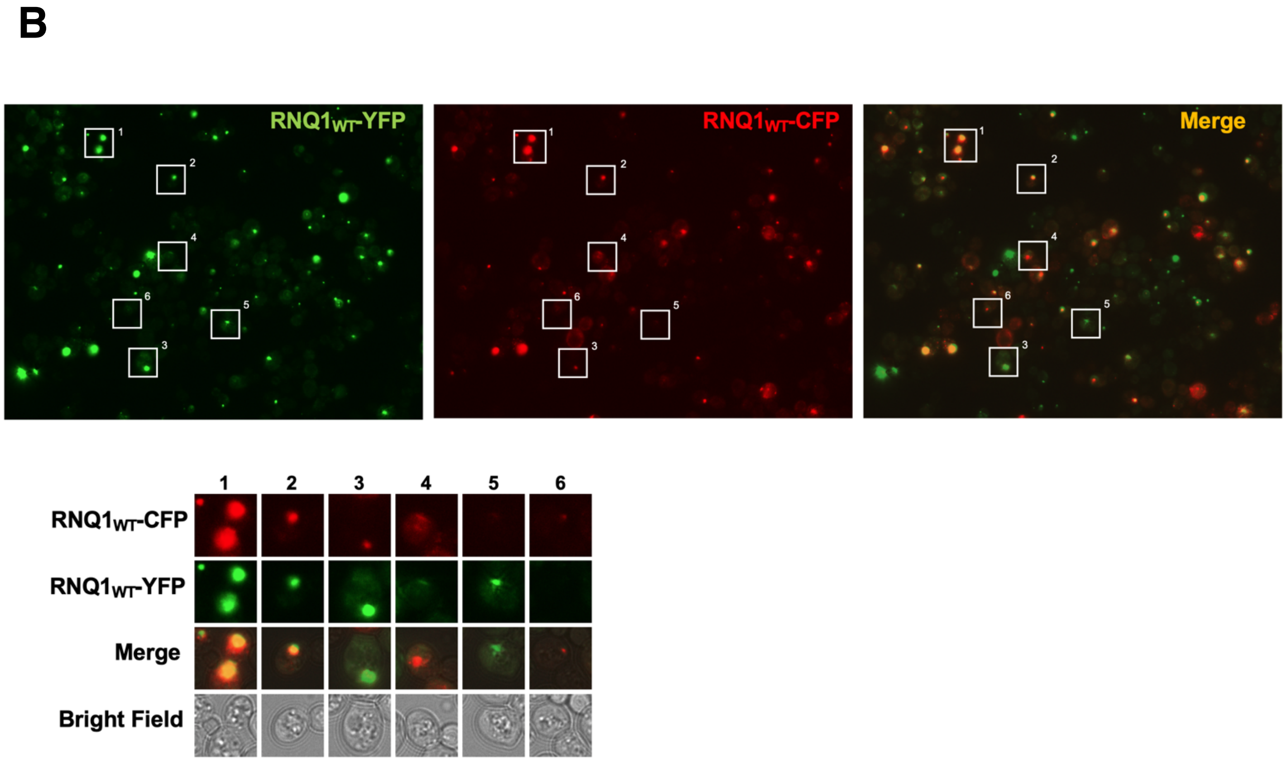

### Supplementary Figure 3

#### Figure S3

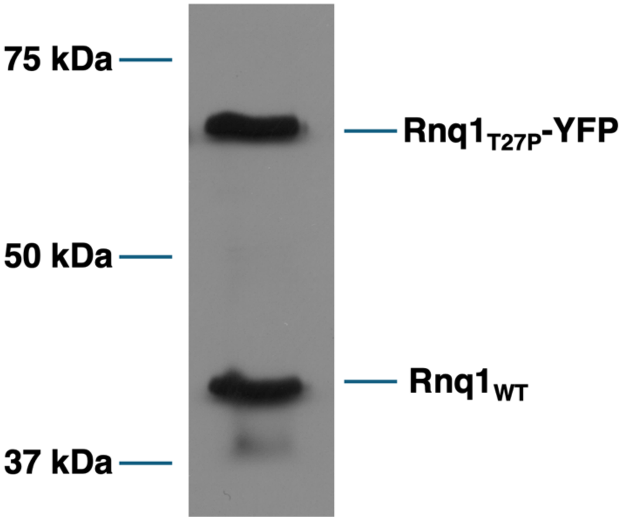
