## Supplemental Table 1 for "Interactions between non-prion and prion domains of Rnq1 direct formation of amyloid vs liquid-like aggregates and create transmission barriers"

**Table S1. Similar arrays of [*PSI*$^+$] variants are induced in cultures expressing Rnq1$_{T27P}$, Rnq1$_{WT}$, or not expressing Rnq1 after shuffling out the *CEN URA3 RNQ1*$_{WT}$ [*PIN*$^+$$_{WT}$] maintainer plasmid.**

| Plasmid after shuffle | [*PSI*$^+$] variants: number isolated (%) | | | | | | | |
|---|---|---|---|---|---|---|---|---|
| | Very strong | Strong | Medium | Weak | Very weak | Leu$^+$ | His$^+$ | Total |
| *RNQ1*$_{WT}$ | 1 (1.0) | 8 (8.6) | 89 (84.8) | 5 (4.8) | 1 (1.0) | 39 (37.1) | 28 (26.7) | 105 (100) |
| *RNQ1*$_{T27P}$ | 1 (1.5) | 1 (1.5) | 62 (91.2) | 4 (5.9) | 0 (0) | 38 (55.9) | 34 (50.0) | 68 (100) |
| EV | 0 (0) | 0 (0) | 37 (100) | 0 (0) | 0 (0) | 15 (40.6) | 19 (51.3) | 37 (100) |

See Results and Materials and Methods for the description of the experiment. [*PSI*$^+$]s isolated from at least eight independent transformants carrying the indicated plasmids were analyzed by the color test on YPD, growth on adenineless media and curability by GuHCl. Results of all three assays were consistent for ~99% [*PSI*$^+$] variants.
