## Supplemental Materials and Methods for "Interactions between non-prion and prion domains of Rnq1 direct formation of amyloid vs liquid-like aggregates and create transmission barriers"

### File S1. Supplementary Materials and Methods

#### Analysis of the *de novo* induction of [$PIN^{+}_{T27P}$]

For the experiment in Figure 3A, the pID193 [*pin*⁻][*psi*⁻] *rnq1-Δ* strain already carrying the *HIS3 GAL-SUP35NM-YFP* [$PSI^{+}$] inducing construct was transformed with the *URA3*, *CEN CUP-RNQ1*$_{T27P}$ plasmid (pID337). Control samples were transformed with *URA3*, *CEN CUP-RNQ1*$_{WT}$ (pID336) or with the pRS416 empty vector. Transformants were selected on and patched onto the SD-UraHis media, then passaged once or twice on SGal-UraHis+50μM $CuSO_4$ (or on control media where neither or only one of the two constructs present in the cell was overexpressed: SD-UraHis, SD-UraHis+50μM $CuSO_4$, and SGal-UraHis). Then cultures were transferred to adenineless media for [$PSI^{+}$] detection (see Materials and Methods for scoring for [$PSI^{+}$]).

In the experiment described in Figure 3B, the pID193 [*pin*⁻][*psi*⁻] *rnq1-Δ* strain already carrying the *HIS3 GAL-SUP3NM-YFP* [$PSI^{+}$] inducing construct was co-transformed with *URA3 CEN CUP-RNQ1*$_{T27P}$ (pID337) and the *CEN LEU2* plasmid carrying *RNQ1*$_{T27P}$ under its original promoter (pID335). Control samples were co-transformed with either *URA3 CEN CUP-RNQ1*$_{WT}$ (pID336) and the *CEN LEU2 RNQ1*$_{WT}$ (pID129), or with the pRS416 and pRS415 empty vectors. Transformants were selected on and patched onto SD-UraLeuHis media and then passaged once or twice on SD-UraLeuHis+50μM $CuSO_4$ (or on control media without copper). Then, yeast were transferred to SGal-UraLeuHis without copper either directly (1 passage in Figure 3B), or after additional passages on SD-UraLeuHis without copper (2, 3, and 4 passages in Figure 3B). Finally, yeast were transferred from SGal-UraLeuHis to adenineless media for [$PSI^{+}$] detection. Additional controls included co-overexpression of Rnq1 and Sup35NM-YFP constructs on SD-UraLeuHis+50μM $CuSO_4$ (0 passages in Figure 3B), as well as all variants of the experiment corresponding to 0, 1, 2, 3, and 4 passages where expression of the $Rnq1_{T27P(or\ WT)}$ and/or Sup35NM-YFP was never induced.

**Preparation of Yeast Cell Lysates for Western Blot Analysis of $Rnq1_{T27P}$ expression**

Yeast cell lysates were prepared essentially as described in Liebman et al. (2006) and Kadnar et al. (2010). Specifically, 50 ml yeast cultures were grown in YPD to mid-log phase. Cells were harvested by centrifugation, washed twice with cold water, and re-suspended in 300μl of protein extraction buffer (50 mM TrisHCl, pH 7.5; 50mM KCl; 10 mM $MgCl_2$; 5% w/v glycerol) supplemented with an antiprotease cocktail for yeast (Sigma; 1:100) and additional 10 mM PMSF. Suspensions were transferred to Eppendorf tubes containing 300μl of 0.5 mm glass beads. Cells were lysed by vortexing at high speed (10 times for 25 s; in between on ice for 25 s). Crude lysates were cleared by low-speed centrifugation (1 min, 600 x g, $4^0$C).

**Recombinant protein purification from *E. coli***

Recombinant protein purification was essentially as described in Kadnar et al. (2010). Specifically, the pJC45-based $Rnq1_{WT}$, $Rnq1_{T27P}$, $\Delta B1C2D3_{WT}$, and $\Delta B1C2D3_{T27P}$ expression constructs were introduced into *E. coli* BL21-AI One Shot cells (Invitrogen). Transformants were grown in LB medium supplemented with 100 μg/ml ampicillin at $37^0$C until mid-log phase ($OD_{600}$=0.4) when protein expression was induced by 1mM IPTG and 0.2% L-arabinose for 1.5 hrs. Cells were harvested by centrifugation ($4^0$C, 1600 x g) and lysed by gentle agitation in lysis buffer (100mM $NaH_2PO_4$, pH 7.4; 350mM NaCl; 8M urea) for 1hr at room temperature (RT). Cell debris was removed by centrifugation at 20,000 x g for 10 min. To bind the 10xHIS-tagged Rnq1, supernatants were incubated for 2 hrs with Ni-NTA Sepharose (Qiagen) equilibrated with the lysis buffer. The slurries were transferred to columns and washed extensively with the lysis buffer. Rnq1 was eluted with the lysis buffer containing 250mM imidazole. UV absorption at 280nm was used to identify protein containing fractions. Centricons (Millipore) were used to concentrate proteins. Protein concentration was determined by the BCA™ Protein Assay Kit (Pierce). Purity of recombinant proteins was estimated by SDS-PAGE as >95%.

**Staining amyloid aggregates in yeast cells with thioflavin T (ThT)**

Staining with ThT was done essentially as described in Douglas et al. (2008). Specifically, cultures of yeast transformants carrying a *CUP-RNQ1*$_{WT}$*-YFP* or a *CUP-RNQ1*$_{T27P}$*-YFP* plasmid in the indicated strain were grown in liquid SD-Ura to $OD_{595}$ ~0.8. Then $CuSO_4$ was added to 50μM and growth was continued for four more hrs. Cells were harvested, resuspended in 1ml of the fixation buffer (50mM $K_3PO_4$, pH 6.5; 1mM $MgCl_2$, 4% formaldehyde) and incubated at RT for 1hr. Cells were washed twice in the PM buffer (0.1M $K_3PO_4$, pH 7.5; 1mM $MgCl_2$), resuspended in 100μl of the PMST buffer (0.1M $K_3PO_4$, pH 7.5; 1mM $MgCl_2$, 1M Sorbitol, 0.1% Tween20) with added 0.6μl of β-mercaptoethanol and 1mg/ml zymolyase 20T (Amsbio), and incubated at 30$^0$C for 15 min. Spheroplasts were washed with PMST, incubated in PMST with 0.005% ThT (Sigma; 20 min, RT, in the dark), and washed 3-4 times with PMST. The CFP filter was used for ThT fluorescence analysis.
