## Supplemental Table 2 for "Interactions between non-prion and prion domains of Rnq1 direct formation of amyloid vs liquid-like aggregates and create transmission barriers"

**Table S2. PCR primers used in this study**

| Number | Name | Sequence |
|---|---|---|
| 7 | *RNQ1* start F | 5’ GGCGGATCC**ATG**GATACGGATAAGTTAATCTC |
| 8 | *RNQ1* stop R | 5’ AGGCCGCGGGTAGCGGTTCTGGTTGC |
| 17 | *RNQ1 ORF* F | 5’ GGTTCCTCCTTTGGAGGA |
| 18 | *RNQ1 ORF* R | 5’ GCTCCAAAGGAGGAAACC |
| 19 | *RNQ1* promoter F | 5’ CCGGAATTCCATGGTATTTCAAACGCA |
| 30 | *RNQ1* terminator R | 5’ TCCGAGCTCAAAATTGTCTTGCAGCCCA |
| 118 | *RNQ1* seq F | 5’ AACCCTTAGAGTCTGCC |
| 119 | *RNQ1* PD seq F | 5’ AGGTCAAGGTCAAGGT |
| 124 | *Bgl*II F | 5’ AACGTATAGCAAAGATCTGGTCC**ATG** |
| 127 | *Bcl*I F | 5’ CGCGTGATCAAAGATCT*GGATCC***ATG** |
| 129 | *Hin*dIII R | 5’ CCCAAGCTTGACTGAGCTGCGGATGT |
| 360 | T27P mut F | 5’ CAGAAGCTGTTGCGAAGTTGCCATCCGCAGCTCAGTCGAAC |
| 361 | T27P mut R | 5’ GTTCGACTGAGCTGCGGATGGCAACTTCGCAACAGCTTCTG |

Sites for restriction endonucleases used for cloning are underlined; the ATG start codon is in bold.
